# *Dillenia indica* fruit extract has Glucose and Cholesterol Lowering effects

**DOI:** 10.1101/804815

**Authors:** Shumsuzzaman Khan, Amrita Bhowmik, SM Badier Rhaman, Siew Hua Gan, Begum Rokeya

## Abstract

**Background:** *Dillenia indica* (*D. indica*) can suppress carbohydrates hydrolysis by inhibiting α-amylase and α-glucosidase. However, there is a lack of understanding of its therapeutic potential as an antidiabetic and anti-hyperlipidemic agent.

**Methods and findings:** Type 2 diabetes (T2D) was induced by a single intraperitoneal injection of Streptozotocin (STZ; 90mg/kg) and hyperlipidemia by feeding with 1% cholesterol, 5% coconut oil and 5% cow fat diet. Administration of *D. indica* extracts in water for four weeks triggered a significant (p≤0.05) reduction in fasting serum glucose (FSG) levels with concomitant improvement in serum insulin levels. Both the water- and ethanol-extract of *D. indica* treated groups showed significant (p≤0.01) reduction in total cholesterol levels by 25% and 19%, respectively. HDL-cholesterol was also augmented (by 14%) in ethanol-extract treated group. Liver glycogen content was higher in the water-extract treated group. Histopathological examination revealed that there was no tubular epithelial cell degeneration or necrosis in the renal tissues or hepatocyte degeneration and sinusoidal dilation in liver tissues in animals that received the water-extract. On the other hand, consumption of *D. indica* extract with 1% cholesterol, 5% coconut oil diet or with a 5% cow fat diet for 14 days significantly reduced serum cholesterol levels in group-lll (60→45 mg/dl; p≥0.05) and -IV (85→66 mg/dl; p≥0.05) hypercholesterolemic model rats. *D. indica* fruit extract also reduced serum TG levels (Group-III: 87→65 mg/dl; Group-IV: 40→90 mg/dl; p≥0.05). Interestingly, treatment with *D. indica* prevented a reduction in serum HDL levels in those hypercholesterolemic model rats. Serum LDL levels were significantly lower in group-III (47→39 mg/dl; p≥0.05) and group-IV (57→44 mg/dl; p≥0.05) hypercholesterolemic model rats after *D. indica* treatment.

**Conclusion:** *D. indica* fruit ameliorates FSG, insulin secretion, glycogen synthesis, and serum lipid profile. Therefore, *D. indica* fruit can be a potential therapeutic agent for diabetic and hyperlipidemia.

**Graphical Abstract:** Extract of *D. indica* in water reduces FSG, serum insulin levels, and ameliorates the serum lipid profile in diabetic model rats without any adverse effects on kidney and liver tissues.

Extract of *D. indica* in ethanol significantly reduces serum cholesterol, TG, LDL with no reduction in HDL levels in hyper-lipidemic model rats.

**Highlights:**

- *D. indica* fruit extracts diminished fasting serum glucose (FSG) levels in STZ-induced type 2 diabetic model rats
- *D. indica* fruit extracts boosted insulin secretion
- *D. indica* fruit extracts showed no toxic effects on the kidney and the liver functions
- Extract in water was more effective in reducing FSG levels than extract in ethanol
- Chronic consumption of 1% cholesterol, 5% coconut oil and 5% cow fat diet was sufficient to make the rat hypercholesterolemic
- *D. indica* fruit extract has the potential to reduce serum cholesterol, TG, LDL with prevention in reduction in serum HDL levels.

## Introduction

Diabetes mellitus is a chronic metabolic disorder characterized by on-going hyperglycemia. Insulin resistance and decreased insulin secretion are the two cardinal features of type 2 diabetes mellitus (T2DM). T2DM is typically diagnosed based on glucose levels, either a fasting plasma glucose (FPG) ≥ 7.0 or a 2-hour 75 g oral glucose tolerance test (OGTT) ≥ 11.1 mmol/L levels. Pre-diabetic classifications include a fasting blood glucose (FBG) of 5.6– 6.9 mmol/L or a 2-hour blood glucose level of ≥7.8 and < 11.1 mmol/L [1]. T2DM is often accompanied by cardiovascular disease, diabetic neuropathy, nephropathy, and retinopathy. An altered lipid profile is common in T2DM patients. Furthermore, reduced hepatic glycogen storage is also observed in diabetes [2–4]. Advancements in the modern lifestyle over the last century have contributed to a dramatic escalation in the incidence of T2DM and hyperlipidemia worldwide [5, 6]. Diabetes mellitus (DM) ranks seventh among the leading causes of death and ranks third when its complications are taken into account [7]. One of the potential complications of T2DM is coronary heart disease due to hyperlipidemia. Hyperlipidemia is a metabolic disorder characterized by successive accumulation of lipids and leukocytes in the arterial wall. This can contribute to many forms of diseases, especially cardiovascular ones such as myocardial infarction and stroke. Moreover, systemic hypercholesterolemia is associated with massive neutrophilia and monocytosis [8–11]. Peripheral leukocyte amount is proportional to the level of cardiovascular complications [12]. Moreover, hypercholesterolemia is positively associated with systemic neutrophil as well as monocyte expansion, suggesting that abnormalities in circulating lipids can influence myeloid cell expansion [13]. Thus the pathophysiological mechanism underlying hypercholesterolemia is the enrichment and accumulation of systemic neutrophils and monocytes which subsequently increase atherosclerosis. Atherosclerosis due to hyperlipidemia (elevated levels of cholesterol, TG, LDL) is the principal cause of mortality affecting people worldwide [14]. According to the World Health Organization (1999), it was estimated that high cholesterol level causes around one-third of all cardiovascular disease worldwide and that there are 10,000,000 people with familial hypercholesterolemia worldwide [15, 16]. Thus, overall prevention and amelioration of T2DM and hyperlipidemia require an integrated, international approach to combat the rapid increase in the number of patients in the forthcoming years [5, 17].

Dietary cholesterol has a direct effect on plasma levels of cholesterol [18–20]. Dietary cholesterol increases serum cholesterol in all common species if high enough loads are given. The extent of increase depends on the compensatory mechanisms such as enhanced excretion of bile acids and neutral sterols and regulation of cholesterol synthesis and absorption [21]. Lipid-lowering agents have been responsible for a 30% reduction of cardiovascular diseases, therefore validating the search for new therapeutic drugs to reduce hyperlipidemia [22]. In addition, natural products have the potential to prevent T2DM or to keep the disease under control [23–25]. The World Health Organization (WHO) has also recommended evaluating the effectiveness of natural products when there is a lack of safe modern drugs [26, 27]. Moreover, it is believed that natural products may have fewer side effects than conventional drugs. One such natural product is *Dillenia indica* (*D. indica*), locally known as chalta or “elephant apple”. The fruit of *D. indica* is a 5-12 cm diameter aggregate of 15 carpels, with each carpel containing five seeds embedded in an edible pulp. *D. indica* fruit is widely used in many indigenous medicinal preparations against several diseases [28] (Supplemental Figure 1).

The enzyme inhibition capacity of the active constituents in *D. indica* has been reported in several studies. For example, betulinic acid in *D. indica* fruits can inhibit tyrosinase [29], and sterols in *D. indica* leaves can inhibit α-amylase and α-glucosidase [30]. Nevertheless, despite its traditional claims [31], limited data is available on the hypoglycemic and anti-hyperlipidemic activities exerted by *D. indica* fruit. Here we hypothesized that *D. indica* fruit may reduce post-prandial hyperglycemia and hyperlipidemia by suppressing carbohydrate hydrolysis and lipid absorption in the gut respectively, which may be beneficial for diabetic and hyperlipidemic control. Thus, the aim of this present study was to investigate the antidiabetic and anti-hyperlipidemic effect of *D. indica* fruit in Streptozotocin (STZ)-induced type 2 diabetic model rats and high fat diet (1% cholesterol, 5% coconut oil and 5% cow fat) induced hyperlipidemic model rats.

Here we showed that oral administration of both *D. indica* extract in water or ethanol in Long-Evans female rats for 28 consecutive days significantly ameliorates serum fasting glucose with concomitant enhancement in insulin secretion and liver glycogen content. Moreover, no noticeable change was observed in the serum creatinine and ALT levels in the extract in water and extract in ethanol treated mice, which is reflected in the histopathological analysis, indicating that the extract is safe for kidney and liver functions. In addition, consumption of *D. indica* extract with 1% cholesterol and 5% coconut oil or a 5% cow fat diet for 14 days significantly reduced serum cholesterol, TG, LDL levels with concomitant enhancement in HDL in the hyperlipidemic model rats.

## Methods

### Plant material

Cultivated matured fruits of *D. indica* (20 kg) were purchased from the local market. Bangladesh National Herbarium ascertained the fruit to be the correct specimen, and a voucher (DACB-35371) was deposited. The fruits were rinsed with fresh sterilized water and sliced into small pieces with a clean knife. *D. indica* slices were naturally dried under the sun. The dried slices (7 kg) were then ground with a blender to yield 1.8 kg of fine powder.

### Preparation of Extract in Ethanol

*D. indica* powder (900 g) was mixed with 5.4 L of 80% ethanol (1:6), before being kept frozen overnight. On a subsequent day, the suspension was filtered with a sterile cloth, followed by filter paper. Approximately 3.3 L of the filtrate was then collected. The main constituents of *D. indica* were extracted by using a BUCHI Rotavapor R-114 followed by incubation in a water bath (55°C) to remove the ethanol. The process yielded approximately 420 ml of filtrate. The semi-dried extract was further dehydrated in a freeze dryer (HETOSICC, Heto Lab Equipment Denmark) at −55°C followed by storage in an amber bottle (−8°C). The dried extract was weighed using a digital balance (GIBERTINI E 42-B). Approximately 150 g (yield: 16.67% w/w) of extract of *D. indica* fruits was produced.

### Preparation of Extract in Water

The *D. indica* fine powder (900 g) was dissolved in 9 L water (1:10) and was kept frozen overnight. The suspension was filtered using sterile cloth. The solution was then re-filtered using filter paper to produce approximately 5.5 L of filtrate. The suspension was dried by a BUCHI Rota vapor R-114 and was incubated in a water bath (70°C) to evaporate the water, yielding approximately 310 mL of filtrate. The semi-dried aqueous extract was further dried in a freeze dryer (HETOSICC, Heto Lab Equipment Denmark) at −55°C. Subsequently, it was stored in an amber bottle (−8°C). Ultimately, 90 g of *D. indica* fruit extract in water was acquired (10% w/w).

### Animal Model for Anti-Diabetic Study

Adult Long-Evans female rats weighing between 170 and 230 g were used. The animals were bred at the Bangladesh Institute of Research and Rehabilitation in Diabetes, Endocrine and Metabolic Disorders (BIRDEM) in the Animal House, Dhaka, Bangladesh. The animals were kept at constant room temperature (22 ± 5°C), with a humidity of 40-70%, and in a natural 12-hour day-night cycle. The rats were sustained with a standard laboratory pellet diet while water was allowed *ad libitum*. All animal procedures were performed according to the National Institute of Health (NIH) guidelines under the protocol as approved by the Institutional Animal Care and Use Committee of BIRDEM.

### Preparation of Type 2 Diabetes Model Rats

STZ (2-deoxy-2-(3-methyl-3-nitrosurea) 1-d-glucopyranose) is a potent alkylating agent that is highly genotoxic and responsible for generating mutations in the cells. STZ contains a highly reactive nitrosurea side chain which is responsible for the initiation of cytotoxic and genotoxic action on pancreatic β-cells. Moreover, STZ is transported into pancreatic β-cells through GLUT2, the glucose transporter, which becomes non-functional in T2DM. Thus, STZ is widely used for the induction of T2DM [32]. Before use, a fresh STZ solution was prepared by dissolving 100 mg STZ in 10 ml (0.1 M) citrate buffer, pH 4-5. The molecular weight for STZ is 265.222, so the molarity for STZ was 3.77 µM. A T2DM condition was created by a single intraperitoneal injection (90 mg/kg) of STZ in rat pups (48 hours old; mean weight 7 g) as previously described [33, 34]. The experiments were done three months after the STZ injection. T2DM can onset at a young age, but most patients are diagnosed at middle age or later. In our study, we focused on adult-onset diabetes by using 3-month-old rats which are an appropriate model for adult-onset. Rats with blood glucose levels of 8–12 mmol/L under fasting conditions were included in the experiments to ensure that STZ had induced type 2 diabetes in all subjects.

### Grouping of Diabetes Model Rats

The rats (n=32) were divided into four groups (n=8). Depending on their group, they were given one treatment per day for 28 consecutive days of one of the following:

1. Type 2 diabetic control group (T2 Control) was given deionized water (10 ml/kg).
2. Type 2 positive control group (Glibenclamide) was treated with glibenclamide (5 mg/10 ml) (9.9 ml H2O + 0.1 ml Twin 20)/kg.
3. Type 2 diabetic extract in water treated group was administered aqueous extract of D. indica (1.25 g/10 ml/kg), which was standardized to 138.89 g fresh fruit of D. indica.
4. Type 2 diabetic extract in ethanol treated group was treated with ethanol extract of D. indica at a dose of 1.25 g/10 ml/kg, which was standardized to 83.34 g fresh fruit of D. indica.

### Animal Model for Anti-hyperlipidemic Study

The rats (n=24) were divided into four dietary groups (I, II, III and IV) of six rats each. The rats of all groups were fed on a standard pellet diet (diet I) and water ad libitum. Experimental diets were supplied each day through a metallic, smooth stomach tube, along with pellet diet. Rats in group-I were fed with lab pellet diet for 24 consecutive days until the end of the experiment. In order to make the rats hyperlipidemic, the rats from groups-II and -III were fed with 1% cholesterol and 5% coconut oil (diet II) for the first ten days of the experiment. Subsequently, group-II was continued on diet II (1% cholesterol and 5% coconut oil) for an additional 14 days. Rats of group-III were fed with 1% cholesterol and 5% coconut oil (for 10 days) in addition to ethanol extract of *D. indica* for the next 14 days. On the other hand, rats from group-IV received a 5% cow fat diet for the first ten days of experiment and subsequently were continued with the feeding of 5% cow fat diet along with ethanol extract of *D. indica* for an additional 14 days.

### Dose for D. indica Extracts

For the oral toxicity study, the Organization for Economic Co-operation and Development (OECD) guideline 425 was followed. Moreover, the histopathological studies on kidney and liver tissues for both extract-treated groups serve as a marker of the safety of *D. indica* extracts. Treatment with the selected doses for the extract of *D. indica* in ethanol at 1.25 g/10 ml/kg did not lead to any mortality; all the animals were found to be alive, healthy and active during the experimental periods.

### Blood Collection Procedure for Biochemical Analysis

The animals were anesthetized using isoflurane before blood collection (400 µl) following amputation of the tail tip. The samples were centrifuged (2500 rpm X 15 minutes), and the serum was transferred to fresh Eppendorf tubes. Serum was refrigerated (−20°C) until further analysis. All the biochemical experiments were performed within two weeks of serum collection.

### Collection and Preservation of Liver and Kidney Tissue for Histology

Following sacrificed by cervical dislocation, the liver and kidney tissues were collected, washed in normal saline, and fixed by using 40% formaldehyde for 24 hours. Subsequently, the tissues were subjected to alcohol dehydration. All tissue samples were washed and embedded by using paraffin and xylene. The tissues were then double-stained.

### Determination of serum glucose levels

Glucose concentration was estimated by glucose oxidase (GOD-PAP, Boheringer Mannheim GmbH) as described previously [35].

### Determination of serum insulin levels

Serum insulin was analyzed by an enzyme-linked immunosorbent assay (ELISA) kit for rat insulin (Crystal Chem Inc., Downers Grove, IL, USA) [36].

### Measurement of Liver Glycogen Content

The glycogen content in the rat liver was measured by using the anthrone-sulfuric acid method as described previously [37].

### Determination of serum ALT and creatinine levels

Alanine aminotransferase (ALT) levels as well as serum creatinine were measured using a Clinical Biochemistry Analyzer (BioMajesty® JCA-BM6010/C).

### Haematoxylin & eosin (H & E) staining

H & E staining was performed as described previously [38]. In order to demonstrate the difference between the nucleus and the cytoplasm, acid (eosin) and basic (haematoxylin) dyes were used. The slides were placed in Harri’s haematoxylin for 10 minutes and rinsed with tap water until the water was colorless. Then they were counterstained with 1% eosin solution for 1 minute. Photomicrographs were acquired by using a transmission microscope (Nikon, Minako, Tokyo) at Dhaka Medical College Hospital, Bangladesh.

### Measurement of serum cholesterol levels

Serum total cholesterol level was measured by an enzymatic colorimetric method (Cholesterol Oxidase /Peroxidase, CHOD-PAP, Randox Laboratories Ltd., UK), as described previously [39].

### Determination of serum TG levels

Serum triglyceride (TG) was measured by an enzymatic colorimetric glycerol-3-phosphate oxidase phenol aminophenazone (GPO-PAP) method (Randox Laboratories Ltd., UK), as described previously [40].

### Determination of serum HDL levels

Serum HDL level was determined by using an enzymatic colorimetric assay (HDL cholesterol E kit, WAKO Diagnostics), as described previously [41].

### Determination of serum LDL levels

Serum LDL level was calculated either indirectly by using the Friedewald Formula [50] or directly by using the Equal LDL Direct Select Cholesterol Reagent as described previously [41]

### Statistical Analysis

Data were analyzed using a Statistical Package for Social Science software version 12 (SPSS Inc., Chicago, Illinois, USA). The data were reported as mean ± SD or as median (range) where appropriate. Statistical analysis was accomplished by using student t-tests (paired and unpaired) or ANOVA (analysis of variance) followed by a Bonferroni post hoc test. A p-value of ≤0.05 was considered statistically significant.

## Results

### D. indica decreased fasting glucose levels

After oral administration of the respective treatments for 28 days, there was a decrease in the fasting serum glucose (FSG) levels of animals in all the groups (Figure 1). Only type 2 diabetic rats treated with extract of *D. indica* in water showed a significant reduction (p≤0.05) in FSG, although the extract in ethanol treated group showed a reduction of FSG level (by 11%) when compared to the baseline (Day 0). As expected, glibenclamide significantly (p≤0.01) ameliorated the diabetic condition on day 28 by a 23% reduction as compared to the baseline (Day 0). These data suggest that *D. indica* fruit can lower fasting serum glucose levels.

**Figure 1.**
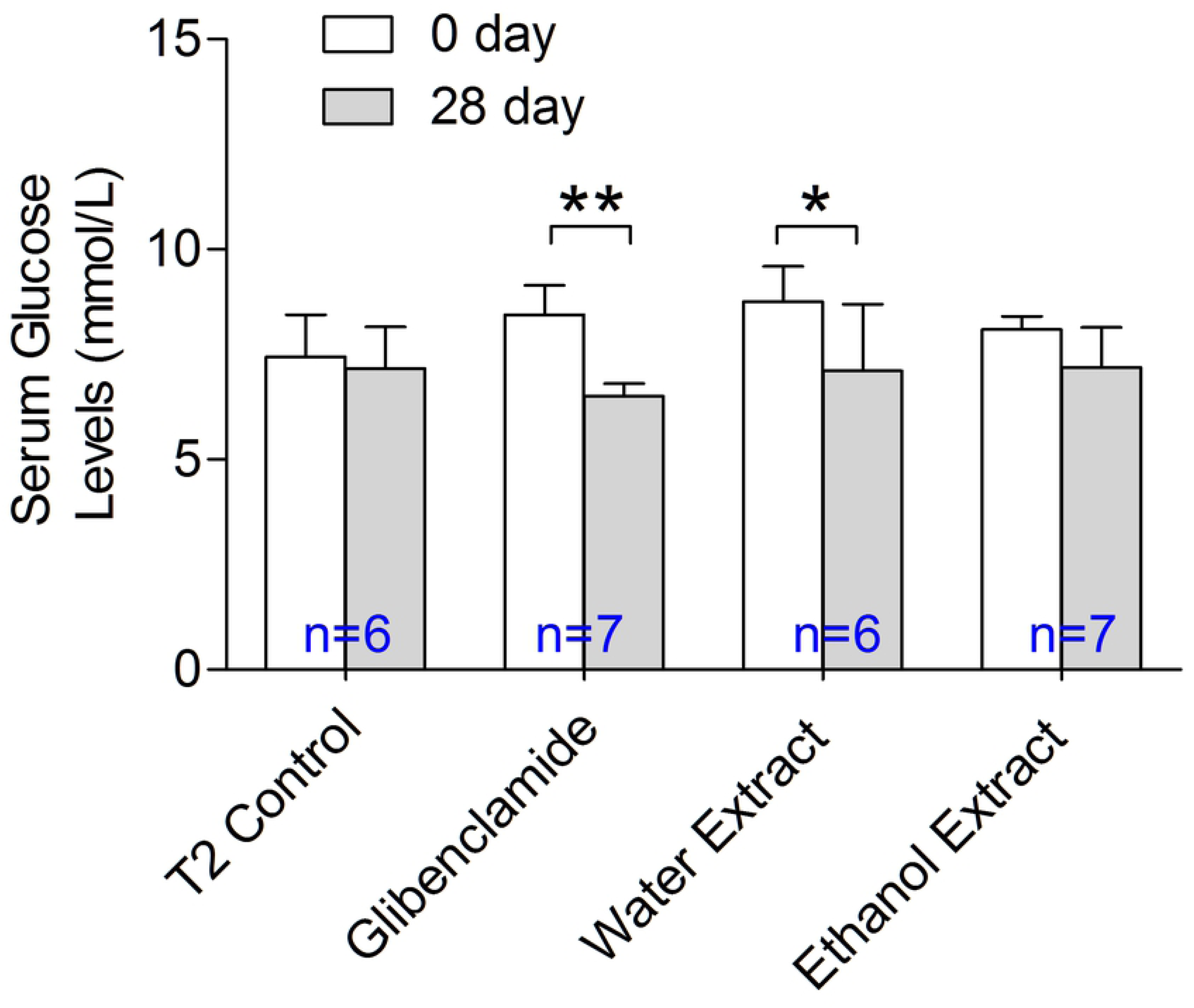
Chronic effect of fruit extracts on fasting serum glucose (FSG) level in STZ-induced type 2 diabetic model rats. Water-extract of *D. indica* significantly decreased FSG levels in diabetic rats. Results are expressed as mean ± standard deviation (SD). Statistical analysis within groups was conducted using a paired t-test while the comparison between groups was done using a one-way ANOVA with post-hoc Bonferroni correction.*p≤0.05; **p≤0.01.

### D. indica increased serum insulin levels

After 28 days, the extract in water treated group showed a significant (p≤0.01) increase (208%) in serum insulin level, while the type 2 control group showed a 44% reduction compared to baseline (Day 0). Moreover, the insulin level was significantly (p≤0.05) higher in the extract in water treated group when compared with the type 2 control group (Figure 2). On the other hand, the glibenclamide treated group showed a 30% increase and the extract in ethanol treated group showed a 19% increase in serum insulin levels compared to baseline (Day 0) (Figure 2). These data indicate that *D. indica* fruit positively modulates pancreatic β-cells to release insulin into the blood.

**Figure 2.**
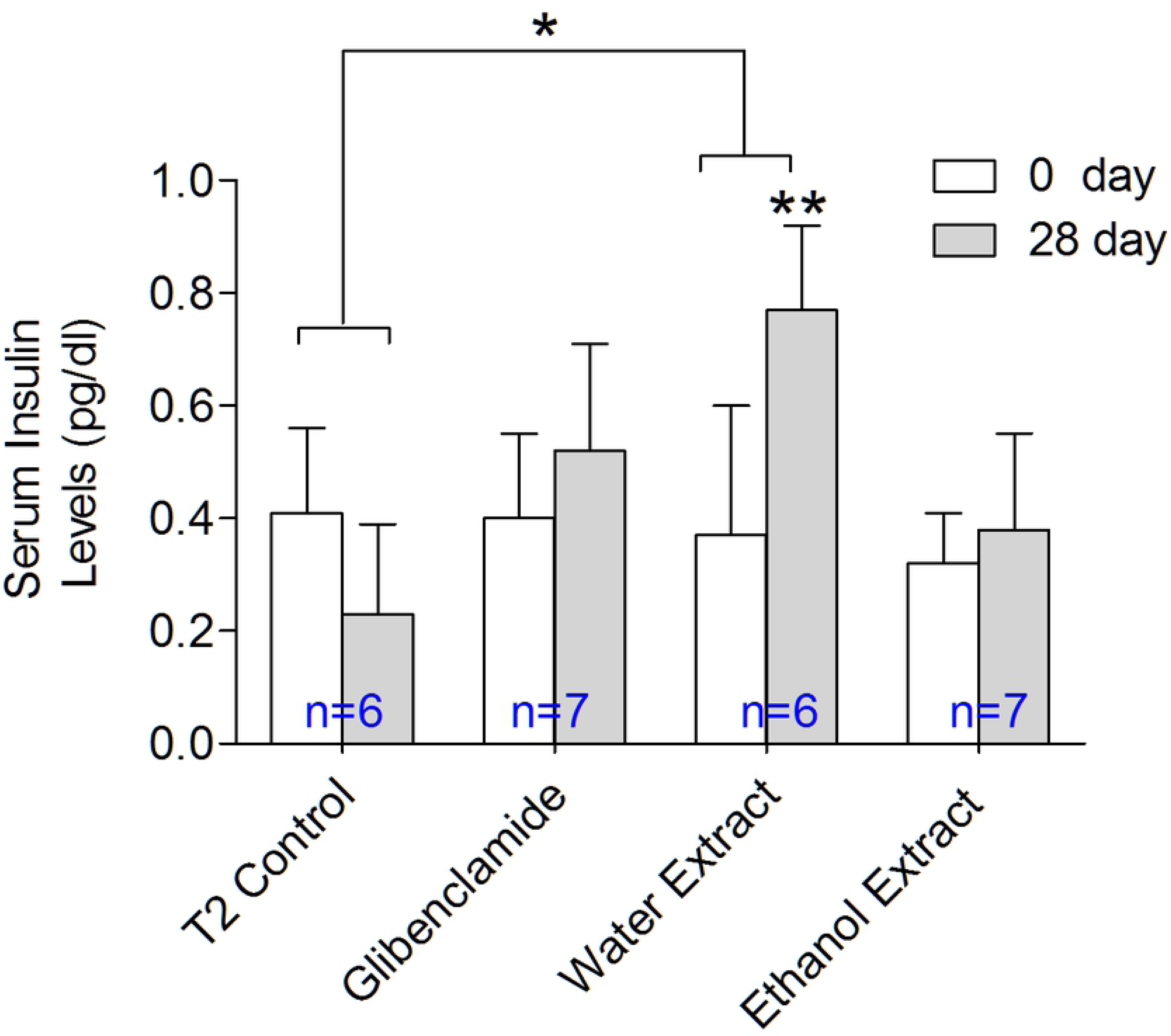
Effect of *D. indica* fruit extracts in serum insulin levels. Water-extract of *D. indica* significantly increased serum insulin levels in STZ-induced type 2 diabetic model rats. Results are expressed as mean ± SD. Statistical analysis within the groups was done using a one-way ANOVA with *post-hoc* Bonferroni correction. *p≤0.05, **p≤0.01.

### D. indica did not affect body weight

The effect of both *D. indica* extracts (in water and ethanol) on the body weight of type 2 diabetic rats during 28 days of chronic administration was observed. The body weight of each rat was taken at seven days’ intervals and was found to increase on average by 2-7% in all groups. However, there was no significant difference among the groups (Figure 3), suggesting that *D. indica* fruit extracts did not affect body weight.

**Figure 3.**
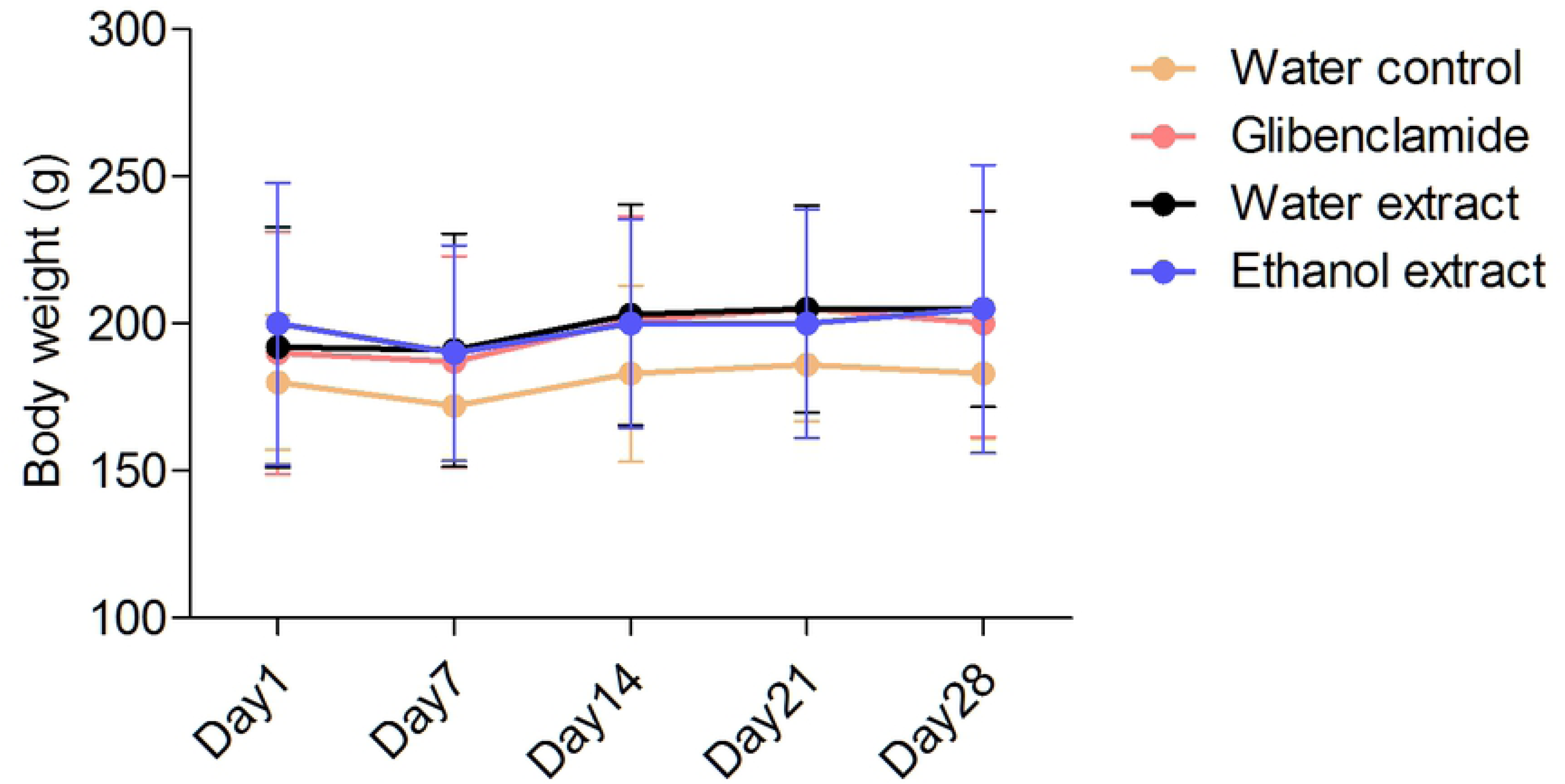
A consequence of *D. indica* fruit extracts on rat body weight. No significant change was observed in rat body weight among the groups after chronic treatment with *D. indica* fruit extracts. Statistical analysis between the groups was done using a one-way ANOVA with post-hoc Bonferroni correction.

### D. indica improved liver glycogen content

The extract in water treated group showed the highest amount (20.39 mg/g) of liver glycogen content among all the groups, whereas liver glycogen content was lowest (7.54 mg/g) in the type 2 control group. There was no significant difference in the liver glycogen content when the glibenclamide and extract-treated groups were compared (Figure 4). Thus, the fruit of *D. indica* enhances liver glycogen content in diabetic rats.

**Figure 4.**
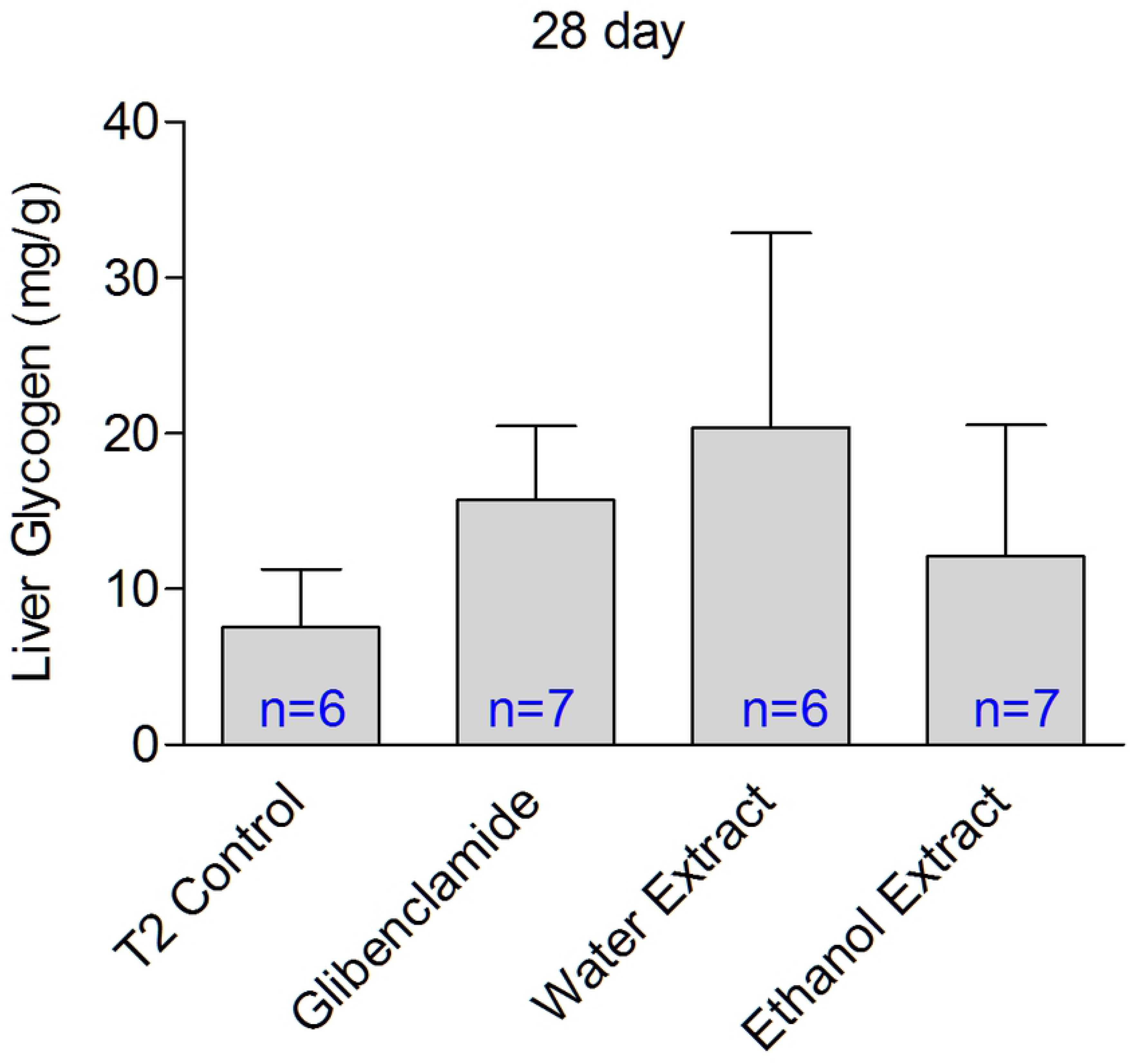
Effect of *D. indica* fruit extracts on liver glycogen content. *D. indica* extract in water showed the highest amount of liver glycogen content among all the groups including the glibenclamide treated group. The results are expressed as mean ± SD.

### Effect of D. indica on serum lipid profile in diabetic model rats

*D. indica* extract in water significantly decreased (p≤0.01) serum cholesterol on Day 28 [serum cholesterol (mean ± SD) mg/dl: Day 0 (75.00 ± 6.37) vs Day 28 (56.00 ± 4.84)] when compared with the glibenclamide-treated group (Figure 5A). Extract of *D. indica* in ethanol caused a significant (p≤0.01) reduction in the total cholesterol level on day 28 [serum cholesterol (mean ± SD) mg/dl: Day 0 (75.00 ± 6.30) vs Day 28 (61.00 ± 2.78)] when compared with type 2 control (Figure 5A). Both the extract in water and the extract in ethanol treated groups showed a reduction in serum TG by Day 28, about 29% (62.00 to 44.00 mg/dl; p≤0.05) and 32% (57.00 to 39.00 mg/dl; p≤0.05), respectively (Figure 5B).

**Figure 5.**
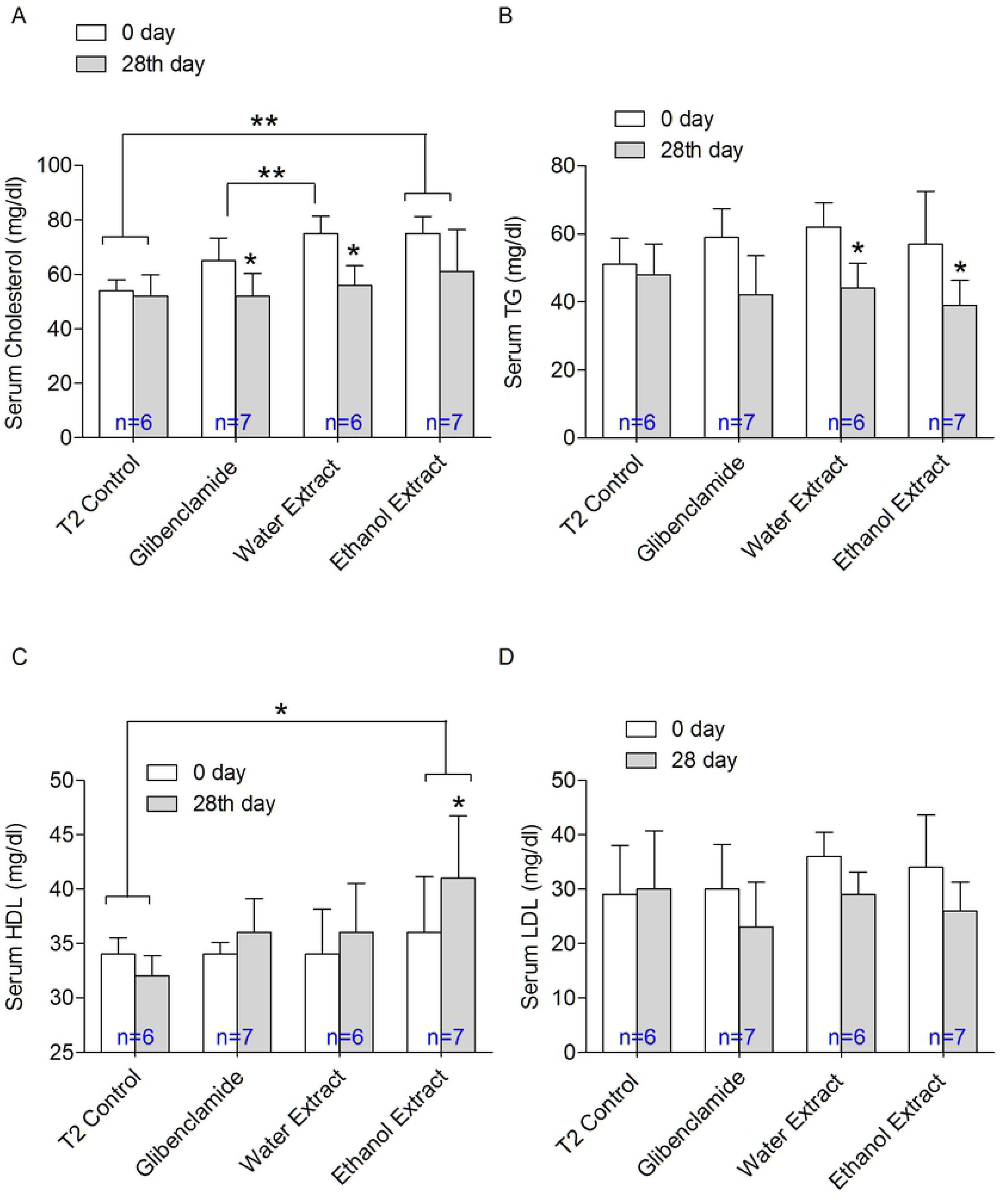
Effect of *D. indica* on serum lipid profile. (A) Both extract in water and in ethanol significantly decreased serum cholesterol levels. (B) Both extracts in water and in ethanol showed a significant reduction in serum TG levels. (C) Extract in ethanol significantly increased serum HDL levels. (D) Reduction in serum LDL levels was also observed by *D. indica* fruit extracts but was not statistically significant. Results are expressed as mean ± SD. Statistical analysis within the groups was done using a one-way ANOVA with *post-hoc* Bonferroni correction. *p≤0.05, **p≤0.01.

The extract of *D. indica* in ethanol significantly increased serum HDL by 14% (36 to 41 mg/dl; p≤0.05), while it reduced LDL by 24% (34 to 26 mg/dl) by Day 28 as compared to the baseline (Day 0). In addition, enhanced HDL cholesterol in the extract in ethanol treated group was statistically significant (p≤0.05) when compared with the type 2 control group. Moreover, both the glibenclamide and extract in water treated groups showed an increase in serum HDL levels by 6% (34 to 36 mg/dl) (Figure 5C). Atherogenic LDL-cholesterol levels were decreased by 24% (34 to 26 mg/dl), 19% (36 to 29 mg/dl), and 23% (30 to 23 mg/dl) for the extract in ethanol, extract in water, and glibenclamide treated groups, respectively (Figure 5D). These data suggest that *D. indica* fruit can ameliorate the lipid profile in diabetic rats.

### Effects of D. indica on serum ALT and creatinine levels

The effects of the extract of *D. indica* in water or in ethanol on kidney and liver functions were also investigated. There was an increase in the serum creatinine levels only in the type 2 control group, although serum creatinine was decreased in glibenclamide (9%), extract in water (13%), and extract in ethanol (6%) treated groups (Figure 6A). In addition, there was no significant change in serum ALT levels in any of the groups (Figure 6B). These data indicate that *D. indica* extracts are safe for kidney and liver functions.

**Figure 6.**
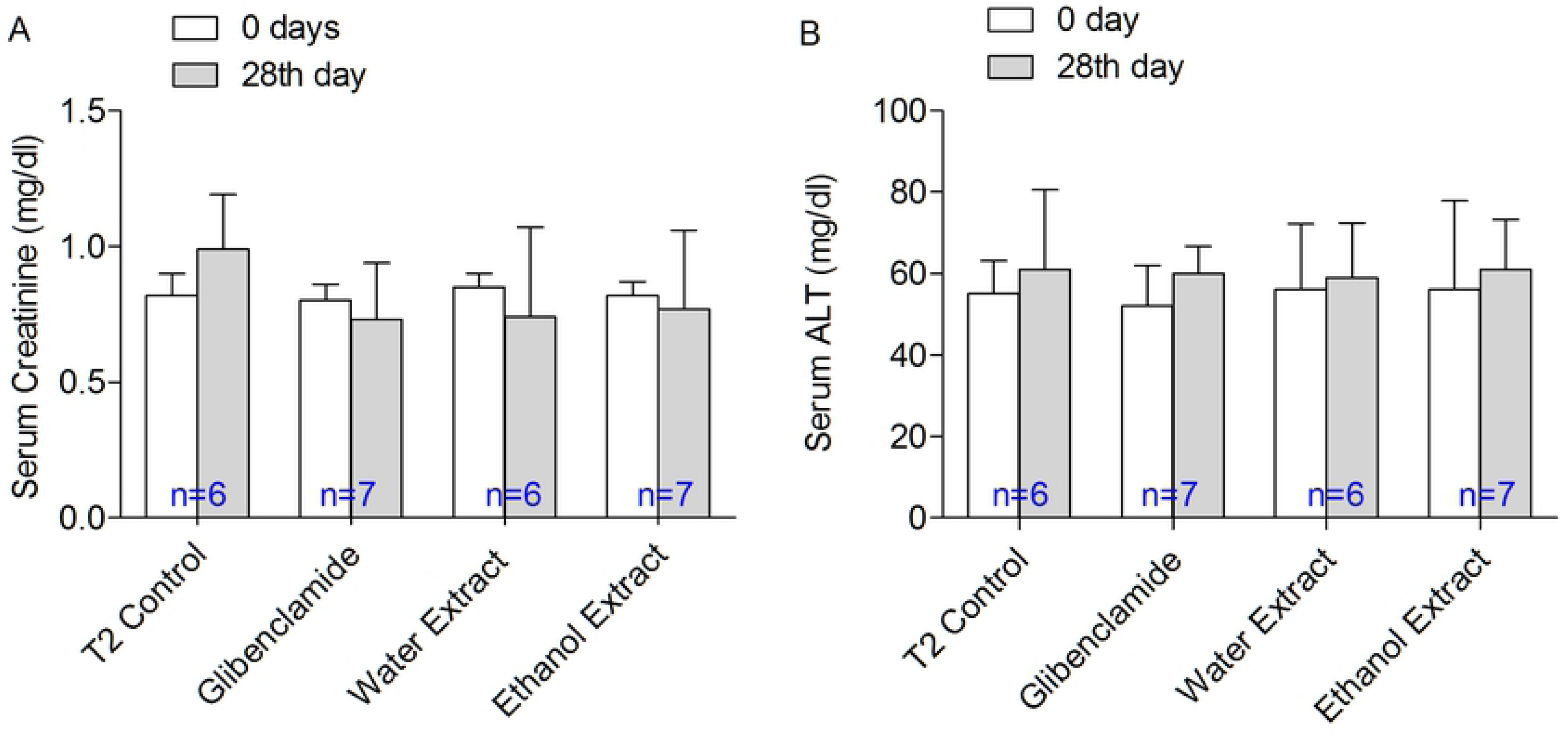
Evaluation of *D. indica* extracts on kidney and liver functions. *D. indica* fruit extracts led to no significant change in the serum creatinine (A) and the serum ALT (B) levels. Results are expressed as mean ± SD. Statistical analysis within the groups was done using a one-way ANOVA with *post-hoc* Bonferroni correction.

### Histopathology

To further confirm the safety of *D. indica* (1.25 g/10 ml/kg) on kidney and liver functions, histopathological examination was performed. Tubular epithelial cell degeneration, necrosis, and hyperemic vessels in the interstitium were examined in the kidney samples. Histological examination revealed that kidney tissues of the type 2 control group were more affected as compared to the extract in water, extract ethanol, and glibenclamide treated groups. Tubular epithelial cell degeneration and tubular epithelial cell necrosis were observed in the kidney tissue of type 2 control group but were absent in the extract in water treated group. The glibenclamide treated group showed well-arranged cells and there was only mild necrosis observed in the extract in ethanol treated group (Figure 7; Table 1), indicating that the aqueous extract of *D. indica* has some renal protective effects.

**Figure 7.**
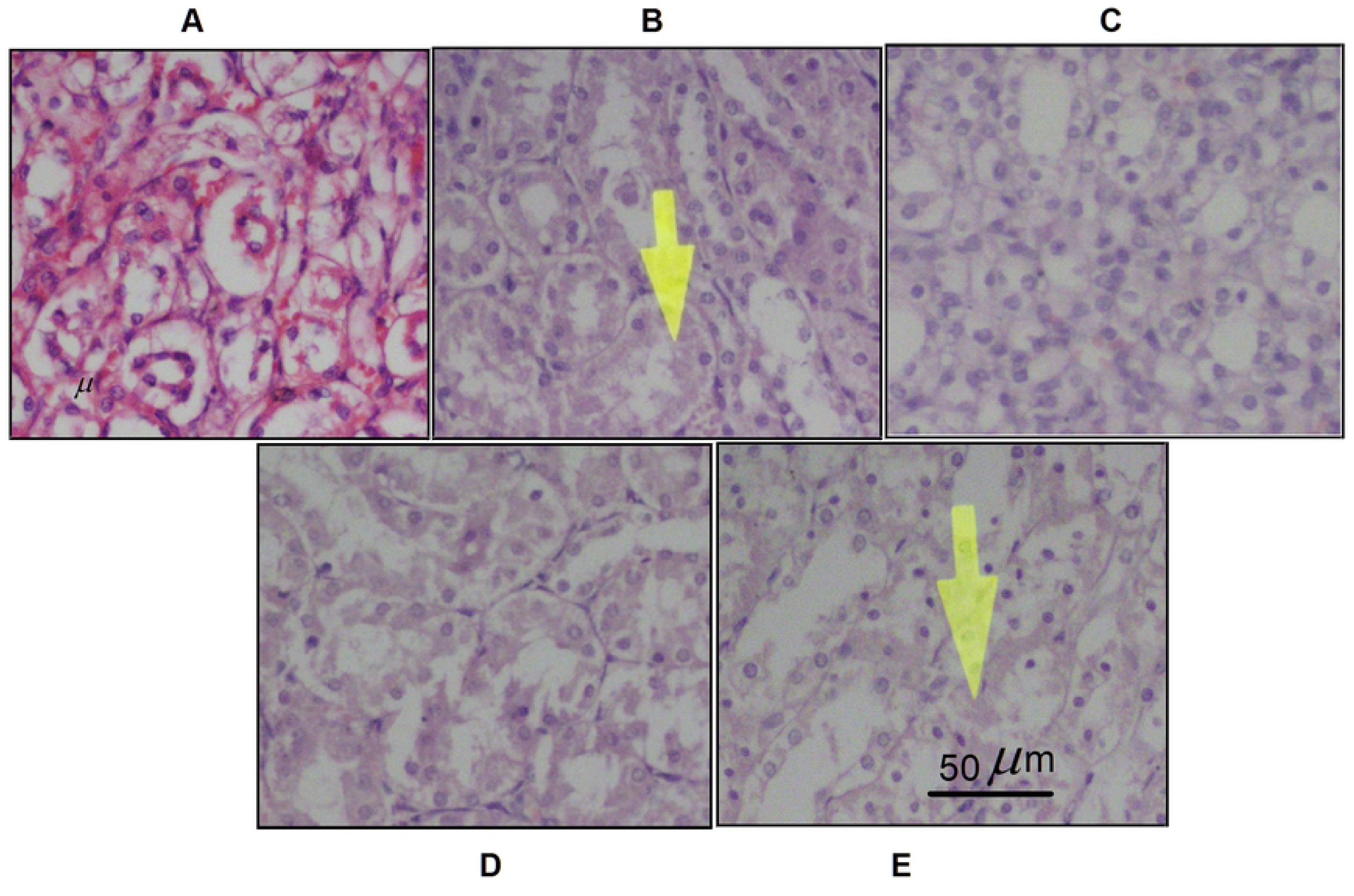
Histological examination of kidney samples after treatment with *D. indica* fruit extracts. (A) The normal control group showed well-arranged kidney cells. (B) The type 2 control group (diabetic) showed mild tubular epithelial cell degeneration and moderate tubular epithelial cell necrosis. (C) The glibenclamide-treated group showed the presence of well-arranged cells. (D) The extract in water treated group showed no toxic effect on kidney cells. (E) The extract in ethanol treated group showed mild necrosis. All figures are observed under a 40x magnification.

**Table 1:** Histopathological changes in kidney samples

| Parameters | Normal group | Type 2 Control group | Glibenclamide group | Water extract group | Ethanol extract group |
| --- | --- | --- | --- | --- | --- |
| Tubular epithelial cell degeneration | - | + | - | - | - |
| Tubular epithelial cell necrosis | - | ++ | - | - | + |
| Hyperemic vessels in the interstitium | - | - | - | - | - |

*\*Histopathologic assessments of the experimental parameters were graded as follows: (-) showing no change and (+), (++) indicating mild and moderate changes respectively.*

In the liver samples, hepatocyte degeneration, sinusoidal dilation, and pleomorphism of the hepatocytes were investigated. Histological examination revealed that the liver tissue of the glibenclamide treated group was more affected as compared to type 2 control, extract in water, and extract in ethanol treated groups. Mild hepatocyte degeneration and sinusoidal dilation were observed in the glibenclamide treated group. Notably, there was no hepatocyte degeneration, sinusoidal dilation, or pleomorphism of the hepatocytes in the extract in water treated group, although mild sinusoidal dilation was observed in the extract in ethanol treated group, suggesting that *D. indica* aqueous extract is safe for liver function (Figure 8; Table 2).

**Figure 8.**
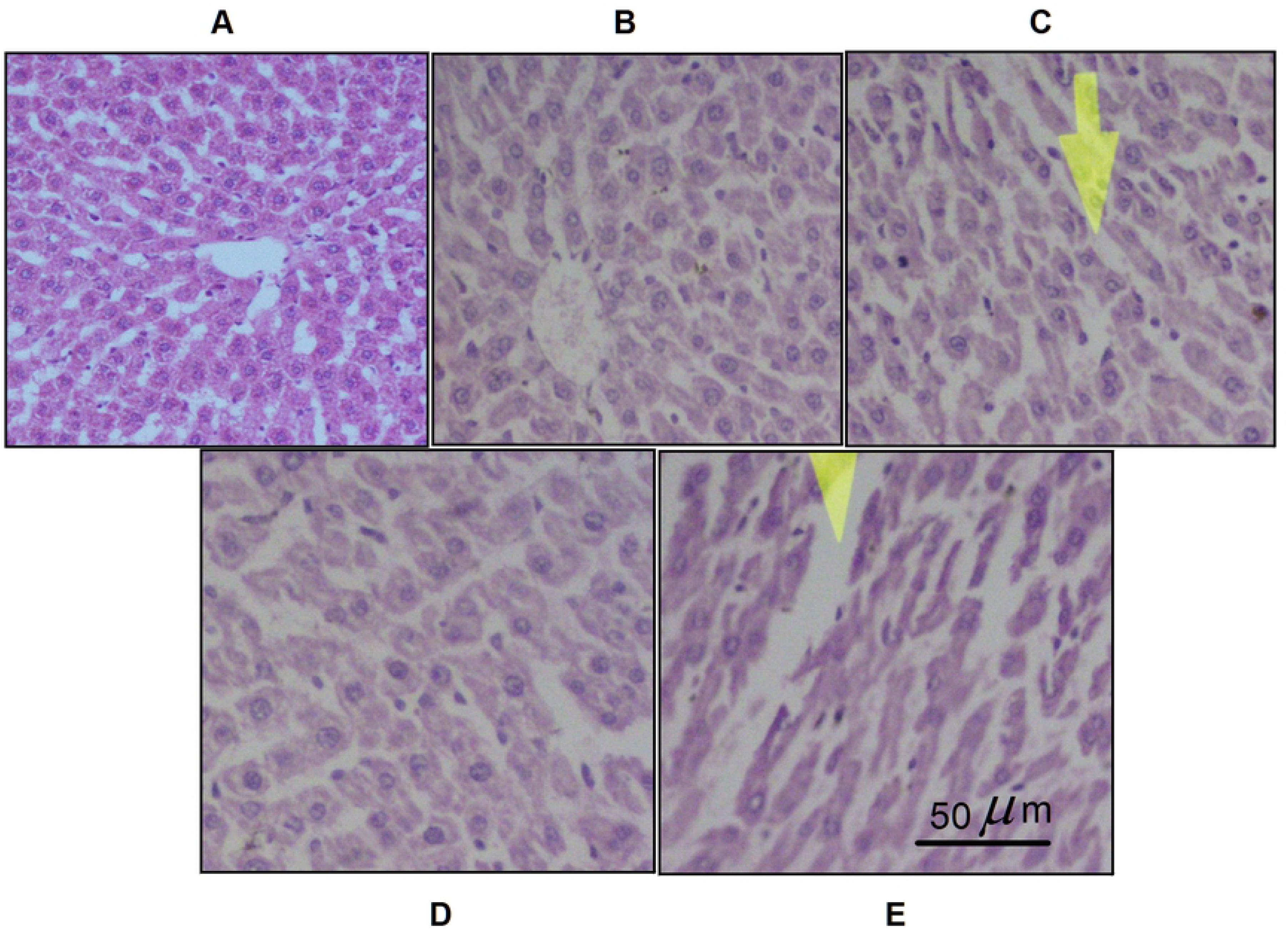
Histological consequences in liver samples after treatment with *D. indica* fruit extracts. (A) The normal control group showed well-arranged liver cells. (B) The type 2 control group (diabetic) showed no significant changes. (C) The glibenclamide-treated group showed mild hepatocyte degeneration and sinusoidal dilation. (D) The extract in water treated group showed no significant pathological changes in liver cells. (E) The extract in ethanol treated group showed mild sinusoidal dilation. All figures are observed under a 40x magnification.

**Table 2:**
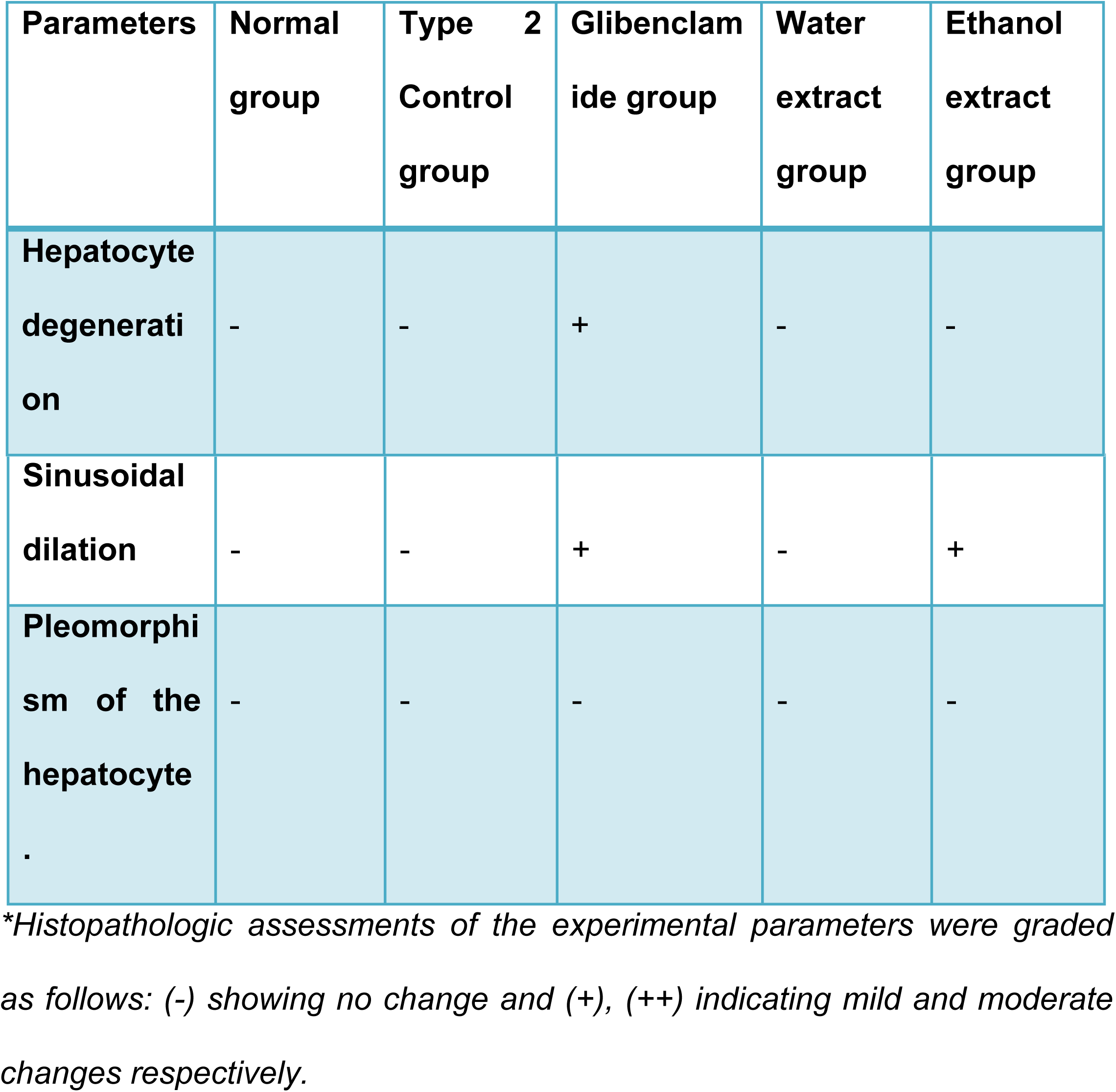
Histopathological changes in liver samples

### Treatment with D. indica fruit extract significantly reduced serum cholesterol levels in hyperlipidemic model rats

Measurement of serum cholesterol is crucial since an altered serum metabolic profile is a potential indicator of many pathological conditions including cardiovascular diseases. In this experiment, we showed that consumption of pellet diet did not enhance serum cholesterol levels in control rats (group I). Moreover, there was also no noticeable change in the control rats among all the groups. In contrast, treatment with 1% cholesterol and 5% coconut oil significantly enhanced serum cholesterol levels at day 10 (41 to 66 mg/dl; p≥0.05), and at day 24 from 66 to 71 mg/dl. Moreover, ANOVA analysis showed that enhancement of serum cholesterol levels in group-ll was significant (p≥0.05) when compared with the normal pellet diet group. These data suggest that consumption of 1% cholesterol and 5% coconut oil has an acute effect on the enhancement of serum cholesterol levels as it was observed at day 10 and further consumption for another 14 days did not lead to any significant enhancement at day 24 (Figure 9).

**Figure 9:**
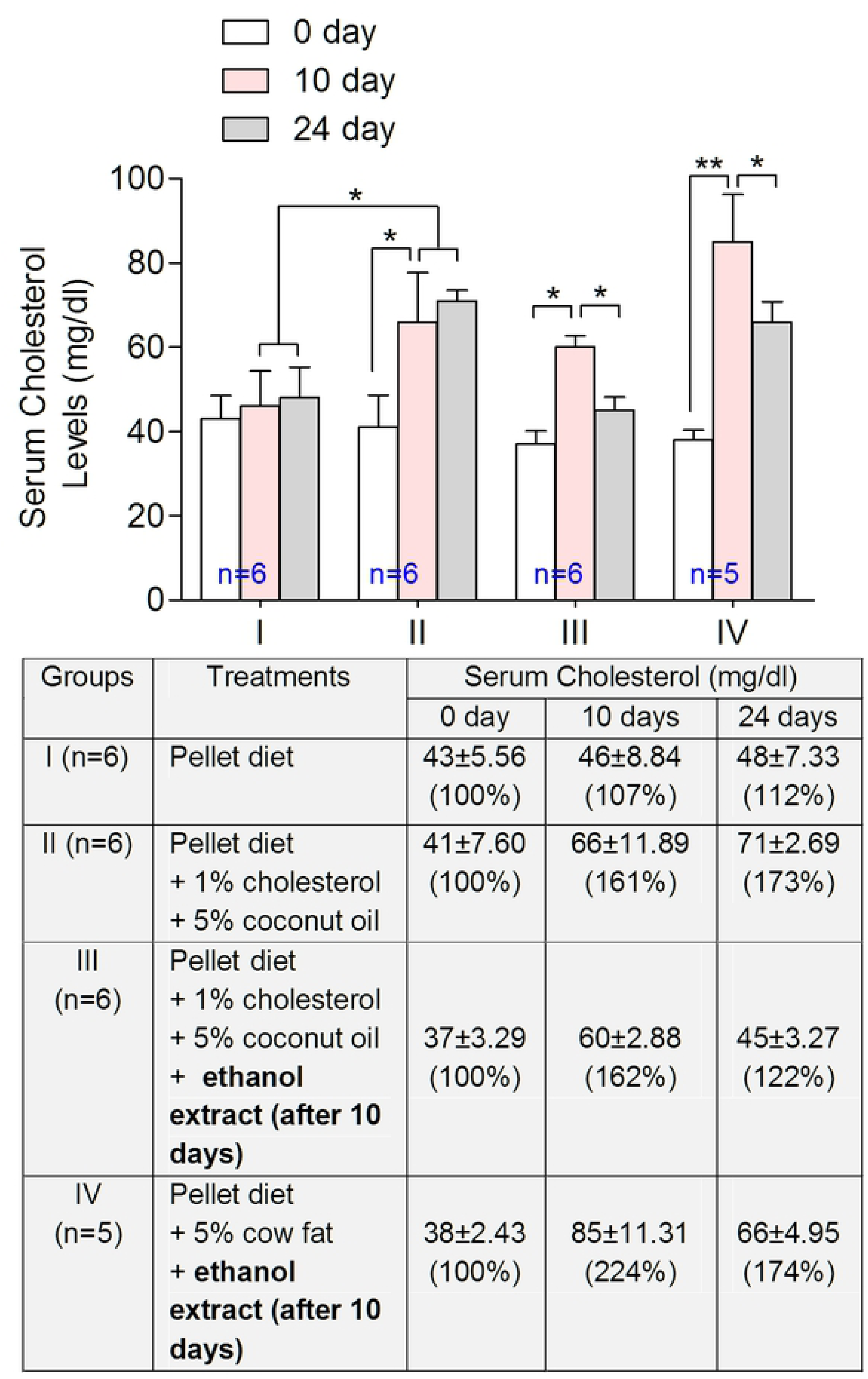
Effect of *D. indica* fruit on serum cholesterol levels in hyperlipidemic model rats. After 14 days treatment with *D. indica* fruit extract in ethanol, significantly reduce serum cholesterol levels in 1% cholesterol and 5% coconut oil diet group and in 5% cow fat treated group. Results are expressed as mean ± standard deviation (SD). Statistical analysis within groups was conducted using a paired t-test while the comparison between the groups was done using a one-way ANOVA with post-hoc Bonferroni correction.*p≤0.05; **p≤0.01.

Interestingly, when *D. indica* fruits extract in ethanol (1.25 mg/kg) was supplemented to the 1% cholesterol and 5% coconut oil treatment in group-lll, those hypercholesterolemic rats showed significant reduction in serum cholesterol levels (60 to 45 mg/dl; p≥0.05), suggesting that *D. indica* fruit extract has potential to reduce serum cholesterol level (Figure 9).

Cow fat (tallow) is primarily made up of triglycerides. In this experiment, a 5% cow fat diet significantly enhanced serum cholesterol levels (38 to 85 mg/dl) in group-lV rats. Moreover, enhancement in serum cholesterol levels was higher (224%) when fed the cow fat diet, than when rats were fed the 1% cholesterol and 5% coconut oil diet (162%), suggesting that cow fat is more potent than 1% cholesterol and 5% coconut oil in inducing serum cholesterol levels. As expected, group-IV rats given the cow fat diet with *D. indica* extract showed a significant reduction (85 to 66 mg/dl; p≥0.05) in cholesterol levels. Although, the reduction was significant, the value (66 mg/dl) was higher than levels (45 mg/dl) in group-III rats (Figure 9). Taken together, these data suggest that extract of *D. indica* in ethanol possesses an anti-cholesterolemic effect with a saturation capacity depending on the fat content and quality of the diet.

### Treatment with D. indica fruit extract significantly reduced serum TG levels in hyperlipidemic model rats

Like was seen with cholesterol levels, the pellet diet did not enhance TG levels in control rats (Group-I). However, treatment with 1% cholesterol and 5% coconut oil for 10 days, enhanced serum TG levels (53 to 80 mg/dl) in group-II rats. Further treatment for an additional 14 days, led to no additional enhancement in serum TG levels, indicating a saturation effect of the extract in lowering TG levels, as was observed with cholesterol levels (Figure 10).

**Figure 10:**
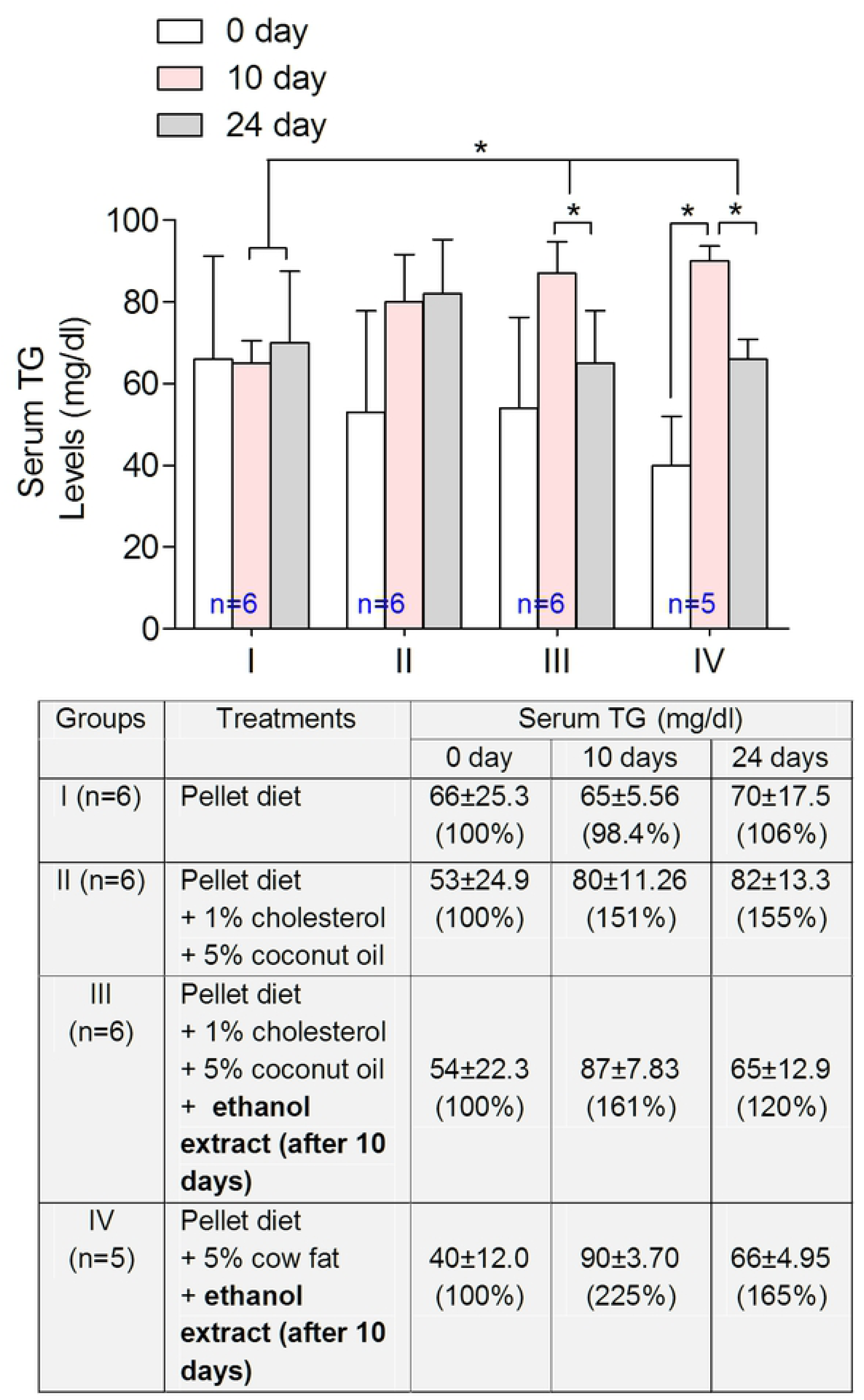
Effect of *D. indica* fruit on serum TG levels in hyper-lipidemic model rats. Consecutive treatment for 14 days with *D. indica* fruit extract significantly reduced serum TG level in the hypercholesterolemia model rats consumed with 1% cholesterol and 5% coconut oil diet group and in 5% cow fat. Results are expressed as mean ± standard deviation (SD). Statistical analysis within groups was conducted using a paired t-test while the comparison between the groups was done using a one-way ANOVA with post-hoc Bonferroni correction.*p≤0.05.

Interestingly, in group-III rats, treatment with 1% cholesterol and 5% coconut oil enhanced serum TG levels at day 10, however, the *D. indica* extract for an additional 14 days significantly reduced serum TG levels (87 to 65 mg/dl; p≥0.05). Treatment with cow fat diet for 10 days significantly enhanced serum TG levels (40 to 90 mg/dl; p≥0.05) in group-IV. Moreover, ANOVA analysis showed that enhancement of serum TG levels in group-II and -III by cow fat diet was statistically significant (p≥0.05) compared with group-I rats. Consecutive treatment with D. indica extract for 14 days with the cow fat diet significantly decreased serum TG levels (90 to 66 mg/dl; p≥0.05). Noticeably, the enhancement in serum TG levels by 1% cholesterol, 5% coconut oil and a 5% cow fat diet in the group-II and group-III rats was statistically significant (p≥0.05) when compared to group-I rats as revealed by ANOVA analysis (Figure 10). Moreover, reduction in serum TG levels in group-III and group-IV rats by D. indica treatment were similar 75% (87 to 65 mg/dl) and 73.3% (90 to 66 mg/dl), suggesting that *D. indica* extract reduced serum TG levels beyond the fat quality and composition of the diet. Taken together, these data suggest that *D. indica* fruit extract in ethanol has a strong serum TG lowering effect in the hyper-cholesterolemic rats beyond the quality of the fat diet.

### Treatment with D. indica fruit extract showed no change in serum HDL levels in hyperlipidemic model rats

In this study, the mean serum HDL level was 38 mg/dl in the pellet diet group (Grpup-l), with no significant change throughout the 24 days. In group-II rats, there no significant change in serum HDL levels for 10 days, however, treatment with 1% cholesterol and 5% coconut oil significantly reduced serum HDL levels (33 mg/dl to 25 mg/dl; p≥0.05) at 24 days, suggesting that the 1% cholesterol and 5% coconut oil diet is sufficient to reduce the amount of HDL in a chronic consumption strategy (24 days) (Figure 11). This was unlike pattern seen with the diets in groups-I and -II which were sufficient to enhance cholesterol and TG within 10 days and plateaued by the end of 24 days. Moreover, ANOVA analysis showed that the reduction of HDL levels in group-II rats was statistically significant (p≥0.05) when compared to group-I (Figure 11).

**Figure 11:**
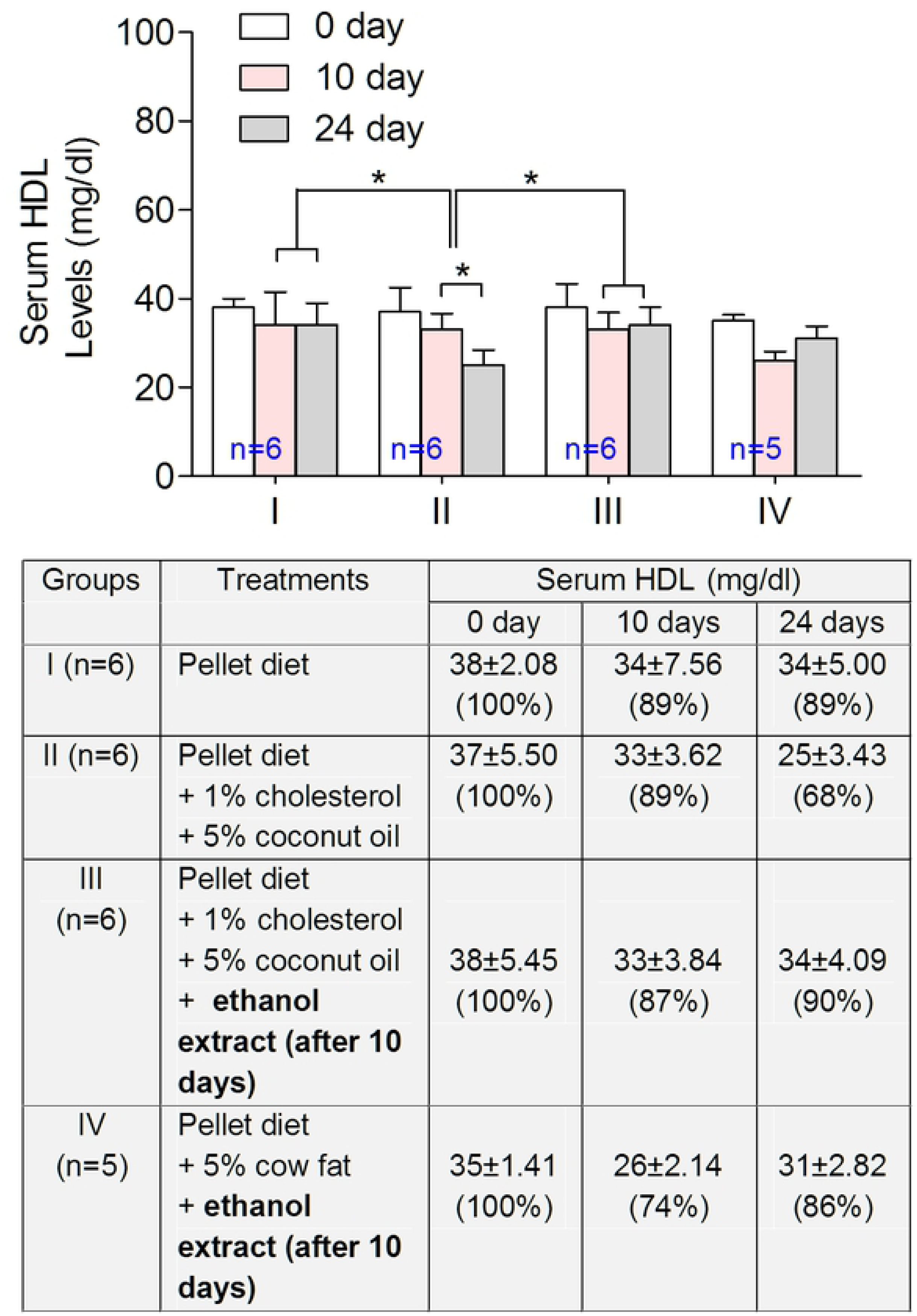
Effect of *D. indica* fruit on serum HDL levels in hyper-lipidemic model rats. *D. indica* fruit extract maintains normal HDL levels in the hypercholesterolemic model rats. 1% cholesterol and 5% coconut oil diet significantly reduced serum HDL levels. Treatment with *D. indica* fruit extract prevented reduction in serum HDL levels in group III and IV rats. Results are expressed as mean ± standard deviation (SD). Statistical analysis within groups was conducted using a paired t-test while the comparison between the groups was done using a one-way ANOVA with post-hoc Bonferroni correction.*p≤0.05.

Interestingly, in group-III rats, treatment with *D. indica* fruit extract with the 1% cholesterol and 5% coconut oil diet for 14 consecutive days suppressed a reduction in serum HDL levels (33 mg/dl to 34 mg/dl) as seen in group-II rats. In group-IV rats, the cow fat diet reduced serum HDL levels by 26% (35 mg/dl to 26 mg/dl) at 10 days, and this reduction was higher than the reduction seen in the 1% cholesterol and 5% coconut oil diet in group-III rats (13%; 38 mg/dl to 33 mg/dl). As expected, treatment with *D. indica* fruit extract with the cow fat diet suppressed the reduction of HDL levels (Figure 11). Taken together, these data suggest that *D. indica* extract has the potential to maintain normal serum HDL levels in hypercholesterolemic rats.

### Treatment with D. indica fruit extract significantly reduced serum LDL levels in hyperlipidemic model rats

The relationship between HDL and LDL is inversely proportional; a high level of HDL is beneficial, while a high level of LDL is bad for health. In this experiment, consumption of the pellet diet for 24 days did not lead to any significant change in serum LDL levels in group-I rats. On the contrary, chronic consumption of 1% cholesterol and 5% coconut oil diet for 24 days in group-II rats significantly enhanced serum LDL levels (27 mg/dl to 42 mg/dl; p≥0.05). Although the 1% cholesterol and 5% coconut oil diet for 10 days were not sufficient for significant enhancement of LDL levels, consumption for 24 days was sufficient to do so (Figure 12).

**Figure 12:**
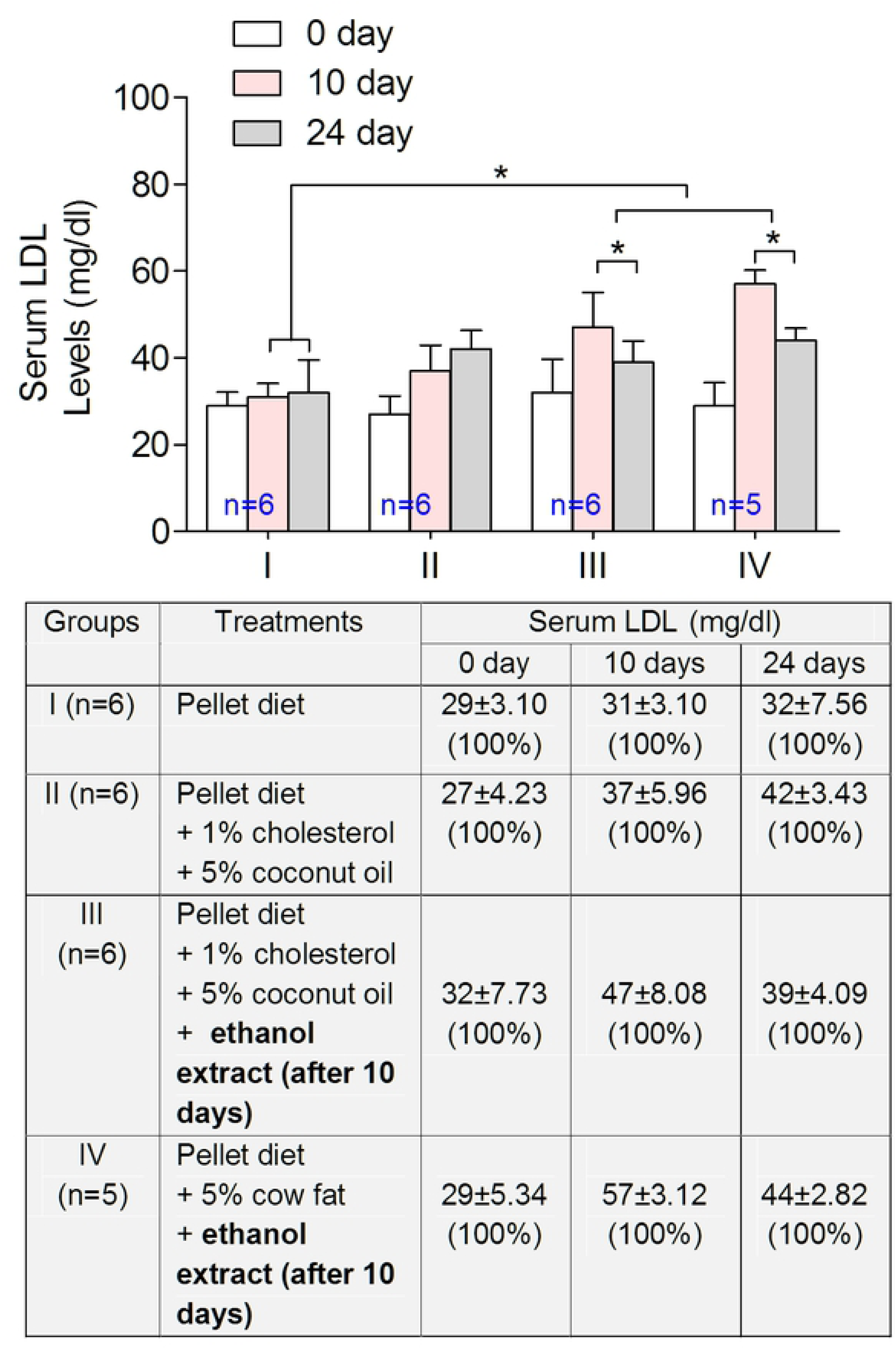
Effect of *D. indica* fruit extract on serum LDL levels in hyperlipidaemic model rats. Chronic consumption of *D. indica* fruit extract for 14 days with 1% cholesterol+5% coconut oil and 5% cow fat diet significantly reduced serum LDL levels in group III and IV rats. Results are expressed as mean ± standard deviation (SD). Statistical analysis within groups was conducted using a paired t-test while the comparison between the groups was done using a one-way ANOVA with post-hoc Bonferroni correction.*p≤0.05.

Interestingly in group-III rats, chronic consumption of 1% cholesterol and 5% coconut oil with *D. indica* fruit extract in ethanol for 14 days significantly lowered serum LDL levels (47 mg/dl to 39 mg/dl; p≥0.05), suggesting that *D. indica* fruit simultaneously reduces cholesterol and LDL levels. As expected, chronic consumption of 5% cow fat diet significantly enhanced serum LDL levels (29 mg/dl to 57 mg/dl, p≥0.05) in the group-IV rats. Moreover, treatment with 5% cow fat diet with *D. indica* fruit extract significantly suppressed serum LDL levels (57 mg/dl to 44 mg/dl; p≥0.05) in group-IV rats (Figure 12). Taken together, these data suggest that a 5% cow fat diet is a potent enhancer of serum LDL levels, while *D. indica* fruit extract ameliorated LDL levels in hypercholesterolemic model rats.

## Discussion

To our knowledge, this is the first study to report on the potential of *D. indica* extracts in ameliorating diabetes and cholesterol levels. In this study, there was a significant reduction of FSG level in the extract of *D. indica* in water and glibenclamide treated groups and a non-significant FSG reduction in the extract in ethanol treated group. Our finding confirms that extracts of *D. indica* in both water and ethanol improved glycemic status in T2DM rats.

Alterations in carbohydrates and lipid metabolism are associated with insulin-resistant states, ultimately causing diabetes [42, 43]. In our study, to determine the probable mechanism(s) of the hypoglycemic effect following chronic treatment with *D. indica* extracts, serum insulin levels of type 2 diabetic rats were measured at baseline on Day 0 and compared to levels on Day 28. Four weeks of treatment with *D. indica* extract in water significantly improved serum insulin levels in type 2 diabetic rats. Thus, it is possible that *D. indica* aqueous extract may enhance cytoplasmic calcium (Ca^2+^) responsible for changes in electrical activity in pancreatic β-cells, leading to enhanced insulin secretion. Moreover, it is plausible that *D. indica* acts on the pancreas to cause a hypoglycemic effect similar to the effects seen with Aloe barbadensis and Litsea glutinosa extracts, both of which ameliorated diabetic conditions [40, 44].

Multiple regulatory mechanisms are involved in glucose homeostasis, and the glucose transporter GLUT2 plays a key role. GLUT2 is expressed in different types of tissues including the liver, kidney, intestine, the pancreatic β-cells, neurons, and astrocytes. Insulin secretion is crucial for GLUT2 activation in pancreatic β-cells, and diabetes causes a significant reduction of GLUT2 expression. A known mutation in GLUT2 is responsible for transient juvenile diabetes [45, 46]. Orexin neurons and GABAergic cells in the CNS express GLUT2 [47]. It has been reported that, in a hypoglycemic condition, closure of K^+^ leak channels and increased activity of AMP-activated protein kinase in GABAergic cells are involved in the activation of GLUT2 to maintain glucose homeostasis [48]. Thus, in contrast, *D. indica* treatment in hyperglycemic conditions may be responsible for the opening of K^+^ leak channels and decreased activity of AMP-activated protein kinase, an alternative mechanism to maintain glucose homeostasis in the CNS.

It has been reported that *D. indica* fruits, which are rich in proanthocyanidins, contain a high amount of B-type procyanidins but a lower amount of B-type prodelphinidins [49]. Proanthocyanidin is a class of polyphenols, which are water-soluble in nature, are found mostly in a variety of fruits. In our study, *D. indica* extract in water conferred stronger glucose-lowering effect than the extract in ethanol. This may be explained by water’s high polarity, leading to an uneven distribution of electron density as compared to ethanol. Moreover, most polyphenols are more soluble in water than in ethanol. However, the extract in ethanol also showed anti-diabetic and significant lipid-lowering activities, so other components present in the extract in ethanol must be playing a role.

At the end of the experimental period, the liver glycogen content was increased in the extract in water treated group. Therefore, it may be hypothesized that the hypoglycemic activity of *D. indica* in T2DM rats occurs due to the increased uptake of glucose for the formation of glycogen due to enhanced glycogenesis. Additionally, it is also possible that suppression of hydrolysis of carbohydrates by inhibiting α-amylase and α-glucosidase may be another underlying mechanism for the antidiabetic effect of *D. indica*, as was seen in other studies [50–52]. Glucagon is a peptide hormone secreted from the pancreatic α-cells [53]. It is elevated under stress conditions and helps to increase energy expenditure [54]. The effect of glucagon is opposite that of insulin, i.e. producing an enhanced glucose level in response to hypoglycemia, which may responsible for the development of diabetes [55]. Thus, *D. indica* fruit extracts may decrease glucagon levels, which may be another possible underlying mechanism for reducing serum glucose levels.

Additionally, T2DM is associated with a marked imbalance in lipid metabolism [56]. Both the extract in water and the extract in ethanol treated groups had a significant (p≤0.01) decrease in serum cholesterol level. There was also a decrease in serum TG (31%) and LDL (24%) levels, while serum HDL cholesterol (14%) was increased in the extract in ethanol treated group. Thus, extract of *D. indica* in ethanol has cholesterol-lowering effects in type 2 diabetic rats, which is comparative with another study [57].

In hyper-cholesterolemic study, consumption of a pellet diet (2.5 Kcal/g) for 24 days resulted in no significant changes in serum cholesterol, TG, HDL and LDL levels (Fig. 1–4; 1^st^ panel), consistent with a previous report that said that mice fed with a pellet diet did not become obese [58], while a high fat diet-induced obesity in rats [59]. Moreover, a hypercaloric pellet diet (6 Kcal/g) led to an increase in body fat, arterial pressure, and high serum glucose, insulin, and leptin, levels, a while normal pellet diet (3.5 Kcal/g) did not lead to an increase in any of these [60]. Thus, we may conclude that the lab pellet diet in this experiment was safe for the rats.

A high-fat diet is key step in making a hypercholesterolemic/hyperlipidemic rat [61]. In our experiment, we used a 1% cholesterol and 5% coconut oil, or a 5% cow fat diet to make the rat hypercholesterolemic. It is well established that dietary cholesterol is a potent enhancer of systemic circulating lipids. Moreover, data from population studies reported that dietary cholesterol is atherogenic beyond LDL concentrations in the blood, although other studies reported that high cholesterol consumption causes moderate increases in serum cholesterol levels [62]. In this experiment, we not only added 1% cholesterol but also 5% coconut oil in the diet because coconut oil normally enhances cholesterol and LDL to a greater extent than cholesterol alone, as reported previously [63]. Moreover, coconut oil decreases myocardial capillary density and aggravates cardiomyopathy [64]. Thus 1% cholesterol with 5% coconut oil diet was sufficient to make the rats hypercholesterolemic in this study. Indeed, chronic consumption of a 1% cholesterol with 5% coconut oil diet for 24 days significantly enhanced serum cholesterol levels and serum TG levels in addition to serum LDL levels. Moreover, we also observed a significant reduction in serum HDL levels in rats after 1% cholesterol with 5% coconut oil treatment (Fig. 1–4; 2^nd^ panel). It has been reported that cows fed with tallow showed higher total cholesterol in plasma than cows fed a low-fat diet [65]. Moreover, 21% tallow with 1.25% cholesterol consumption for six weeks led to renal dysfunction and atherosclerosis in the rats [66]. In addition, a beef tallow diet promoted body fat accumulation and reduced norepinephrine turnover rate in brown adipose tissue by reducing sympathetic activity [67]. Thus, cow fat can be a good atherogenic diet to make rats hypercholesterolemic. In this experiment, a 5% cow fat diet significantly enhanced serum cholesterol and LDL levels after 10 days’ treatment (Fig. 1 and 4; 4^th^ panel). Moreover, 5% cow fat also enhanced serum TG levels (Fig 2; 4^th^ panel). A further extension of the cow fat effect was a reduction in serum HDL levels (Fig. 4; 4^th^ panel). Taken together, 1% cholesterol with 5% coconut oil or a 5% cow fat diet is enough to make the rat hypercholesterolemic.

The results of *D. indica* on hypercholesterolemic model rats showed that there was a significant decrease in serum cholesterol level in the group treated with 1% cholesterol and 5% coconut oil diet and in the group treated with a 5% cow fat diet after 14 days of *D. indica* extract supplementation (Fig. 1; 3^rd^ and 4^th^ panels). Thus *D. indica* fruit possesses therapeutic potential to reduce serum cholesterol levels. The possible mechanisms behind the reduction in serum cholesterol levels may include proanthocyanidin (plant sterols) which is a class of polyphenols found in a variety of fruits including *D. indica* [49]. It is well established that plant sterols are a potent therapeutic target for cardiovascular health [68]. In addition, other components present in the *D. indica* extract besides proanthocyanidins may contribute to the lipid-lowering activity. To maintain lipid homeostasis, cholesterol absorption in the intestine is a significant physiological process [69]. In fact, targeting the cholesterol absorption pathway is one of the most important pharmacological interventions for hyperlipidemia [70]. A carbohydrate diet is positively correlated with serum lipid levels [71]. Chromatographic separation of *D. indica* leaves extract revealed the existence of betulinic acid, quercetin, β sitosterol, and stigmasterol palmitate [72], all of which significantly inhibited the activities of α-amylase and α-glucosidase, which may contribute to the lipid-lowering effects. Phytosterols (Proanthocyanidins) can cross the blood-brain barrier (BBB) [73]. Therefore, it may possible that proanthocyanidins in *D. indica* inhibit HMG-CoA reductase and reduce cholesterol biosynthesis. It has been reported that bioactive palmitoleate can prevent atherosclerosis by suppressing organelle stress and inflammasome activation [74]. Moreover, it has also been reported that phytosterols in *D. indica* possess antioxidant [49] and anti-inflammatory activities by inhibiting the production of tumor necrosis factor-alpha [75] and may be the crucial underlying mechanism to prevent atherosclerosis. Microscopy and phytochemical analysis indicated the presence of xylem fibers, phytosterols, pro-anthocyanidins, terpenoids, glycosides, fatty acids, flavonoids, and phenolic compounds in *D. indica* [49, 76]. *D. indica* is also rich in different types of soluble and insoluble dietary fibers. Phytosterols reduce cholesterol levels by different mechanisms, such as by competing with cholesterol absorption in the gut and in dietary mixed micelles. Additionally, sterols compete with cholesterol for solubilization, co-crystallize with cholesterol to form insoluble mixed crystals and inhibit cholesterol hydrolysis by lipases [77–79]. Therefore, phytosterols in *D. indica* extracts may inhibit cholesterol absorption and hydrolysis in the gut.

Hypercholesterolemic model rats treated with *D. indica* extract for 14 days showed a significant reduction in serum TG levels in the 1% cholesterol and 5% coconut oil or the 5% cow fat diet group (Fig. 2; 3rd and 4th panels). *D. indica* extract may interfere with TG biosynthesis and absorption. Overall, lipase is a critical enzyme to digest dietary TGs. Thus the sterols in *D. indica* fruit extract in ethanol may inhibit lipase, thereby ameliorating fat malabsorption in the intestinal lumen by forming an irreversible bond with fat molecules and ultimately leading to fecal excretion without degradation. Moreover, it has been reported that polyphenols in boysenberry and okra extract suppressed enhancement of TG in plasma [80, 81]; this is consistent with our study, where polyphenol in *D. indica* fruit may lower TG levels in hypercholesterolemic rats. Diacylglycerol acyltransferase 2 (DGAT2) catalyzes the last step of TG biosynthesis in rodents [82]. Thus, we do not exclude the possibility that the *D. indica* extract may inhibit DGAT2 and reduce hepatic TG synthesis.

Chronic consumption of a 1% cholesterol and 5% coconut oil diet or a 5% cow fat diet for 24 days significantly reduced serum HDL levels but treatment with *D. indica* fruit extract suppressed reduction and restored normal HDL levels (Fig. 3). ATP-binding cassette protein A1 (ABCA1), lecithin-cholesterol acyltransferase, and apolipoprotein (apo) A-I are the key components for HDL biosynthesis [83]. ABCA1 exports cholesterol from the membranes to the nascent HDL, while ABCG1 (G Member sub-family of ATP-binding Cassette transporter) transports cholesterol to mature HDL; the key features of cholesterol homeostasis [84]. One possible mechanism to reduce HDL levels with chronic consumption of 1% cholesterol and 5% coconut oil is by downregulation of the genes associated with ABCA1, LCAT, and APOAI molecules. On the contrary, *D. indica* fruit extract could restore and/or up-regulate these gene functions to maintain HDL homeostasis

Both a 1% cholesterol and 5% coconut oil diet, and a 5% cow fat diet significantly enhanced serum LDL levels, and, more importantly, *D. indica* fruit extract significantly reduced serum LDL levels in both groups (Fig. 4). To investigate the underlying mechanism of such reduction, it is important to understand the pathophysiology of LDL. After entering into the artery wall endothelium, LDL initiates atherosclerotic plaque formation by binding to glycosaminoglycans [85]. LDL is oxidized, thus producing modified apoB by altering lysine residues [86]. The modified LDL is recognized by the macrophages and internalized by endocytosis, forming foam cells [87]. These foam cells release cytokines, initiating inflammatory reactions [88]. As a consequence, the cells of the arterial wall proliferate and produce collagen. The plaque enlarges, eventually leading to blood clot formation and blockage of the vessel. Several studies have reported that HDL prevents LDL oxidation, a crucial initial step in LDL pathophysiology, and, thus, reducing LDL levels [89]. In this experiment, *D. indica* fruit extract maintained normal HDL levels in hypercholesterolemic model rats, which may contribute to reducing LDL levels by inhibiting LDL oxidation. In addition, LDL enhances CXCR2 (cytokine) expression, facilitating neutrophils accumulation [90]. On the other hand, monocytes use CCR2 for entry into the atherogenic lesion [91, 92]. Therefore, it may possible that *D. indica* fruit extract may reduce CXCR2 and CCR2 expression and reduce atherosclerosis.

The muscles use creatine to make energy and produce creatinine as a waste product. High levels of creatinine in the blood may indicate diabetic nephropathy. Hyperglycemia enhances the level of reactive oxygen species, which then facilitate the formation of glycation end products as implicated in the pathogenesis of diabetic nephropathy [93, 94]. In our study, treatment with *D. indica* extracts suppressed enhancement of creatinine levels in diabetic rats. On the other hand, ALT is an indicator of liver function. It has been reported that elevated serum levels of ALT are significantly associated with increased diabetic risk [95, 96]. In this study, there was no significant change in serum ALT levels in the *D. indica* extract-treated groups, indicating that *D. indica* does not affect liver function. These observations are correlated with the histological changes as extract of *D. indica* in water showed no tubular epithelial cell degeneration, necrosis, and hyperemic vessels in the interstitium of the kidney tissues and no hepatocyte degeneration, sinusoidal dilation, and pleomorphism of the hepatocytes in liver tissues. Interestingly, the glibenclamide-treated group showed mild hepatocyte degeneration and sinusoidal dilation in our study. Moreover, it has been reported that glibenclamide reduced pro-inflammatory cytokine production and reduced glutathione levels in diabetic patients [97], which be the underlying mechanism of observed hepatocyte degeneration and sinusoidal dilation in our study. Thus, our study not only defines an emerging role of *D. indica* fruit as an anti-diabetic and anti-hyperlipidemic agent but also opens a new window on the safety concerns of glibenclamide treatment.

## Conclusions

*D. indica* fruit has a glucose-lowering effect and enhances insulin secretion as well as glycogen synthesis. It decreases total cholesterol levels and increases HDL-cholesterol. It does not affect renal and liver functions in terms of creatinine and ALT, and, therefore, has the potential to be an anti-diabetic and cholesterol-lowering agent.

## Acknowledgments

The authors are grateful to the technical staff of the Research Division of Bangladesh Institute of Research and Rehabilitation in Diabetes, Endocrine and Metabolic Disorder (BIRDEM) for their assistance. The authors offer thanks to Dr. Zillur Rhaman Khan and Dr. Hemale from Dhaka Medical College for histopathology. The authors are also grateful to Mr. Omrito and Dr. Karim from the Bangladesh Rice Research Institute for their contribution to grinding the sun-dried *D. indica*.

## Author Contributions

S.K and B.R designed the research. S.K performed all the experiments and carried out all the data analysis and figure preparation except ELISA experiment by B.R in Fig. 3. A.B contributed to data analysis (SPSS) and the collection of biological samples (blood, liver, and kidney). S.K; S.H.G; SM.B.R; and B.R wrote and edited the manuscript.

## Edited by

Renee Cockerham, Program Manager, Office of Postdoctoral Scholars, University of Maryland, Baltimore, USA.

## Supplementary Materials

**Supplemental Figure 1:** *D. indica* fruit

## References

1. American Diabetes A. 2. Classification and diagnosis of diabetes: standards of medical care in diabetes-2019. Diabetes care 2019; 42:S13–S28.

2. Lyssenko V, Almgren P, Anevski D, Perfekt R, Lahti K, Nissén M. Predictors of and longitudinal changes in insulin sensitivity and secretion preceding onset of type 2 diabetes. Diabetes 2005; 54(1): 166–74.

3. Wu Y, Ding Y, Tanaka Y, Zhang W. Risk factors contributing to type 2 diabetes and recent advances in the treatment and prevention. Int J Med Sci. 2014; 11(11): 1185–200.

4. Ferrannini E, Lanfranchi A, Rohner JF, Manfredini G, Van-de WG. Influence of long-term diabetes on liver glycogen metabolism in the rat. Metabolism 1990; 39(10): 1082–8.

5. Zimmet P, Alberti KG, Shaw J. Global and societal implications of the diabetes epidemic. Nature 2001; 414:782–787.

6. Franks PW, McCarthy MI. Exposing the exposures responsible for type 2 diabetes and obesity. Science 2016; 354:69–73.

7. Marathe CS, Rayner CK, Bound M, Checklin H, Standfield S, Wishart J, Lange K, Jones KL, Horowitz M. Small intestinal glucose exposure determines the magnitude of the incretin effect in health and type 2 diabetes. Diabetes 2014; 63:2668–2675.

8. Soehnlein O, Swirski FK. Hypercholesterolemia links hematopoiesis with atherosclerosis. Trends Endocrinol Metab. 2013; 24:129–136.

9. Rader DJ, Daugherty A. Translating molecular discoveries into new therapies for atherosclerosis. Nature. 2008; 451:904–913.

10. Lin J, Yang R, Tarr PT, Wu PH, Handschin C, Li S, et al. Hyperlipidemic Effects of Dietary Saturated Fats Mediated through PGC-1β Coactivation of SREBP. Cell. 2005;120:261–273.

11. Hansson GK. Inflammation, atherosclerosis, and coronary artery disease. N Engl J Med. 2005; 352:1685–1695.

12. Friedman GD, Klatsky AL, Siegelaub AB. The leukocyte count as a predictor of myocardial infarction. N Engl J Med. 1974; 290:1275–1278.

13. Rothe G, Gabriel H, Kovacs E, Klucken J, Stohr J, Kindermann W, et al. Peripheral blood mononuclear phagocyte subpopulations as cellular markers in hypercholesterolemia. Arterioscler Thromb Vasc Biol. 1996; 16:1437–1447.

14. Lusis AJ. Atherosclerosis. Nature 2000; 407:233–241.

15. Garcia CK. Autosomal Recessive Hypercholesterolemia Caused by Mutations in a Putative LDL Receptor Adaptor Protein. Science 2001; 292:1394–1398.

16. Abifadel M, Varret M, Rabès JP, Allard D, Ouguerram K, Devillers M, et al. Mutations in PCSK9 cause autosomal dominant hypercholesterolemia. Nat Genet. 2003; 34:154–156.

17. Bohula EA, Scirica BM, Inzucchi SE, McGuire DK, Keech AC, Smith SR, Kanevsky E, Murphy SA, Leiter LA, Dwyer JP, et al. Effect of lorcaserin on prevention and remission of type 2 diabetes in overweight and obese patients (CAMELLIA-TIMI 61): a randomised, placebo-controlled trial. Lancet 2018; 392:2269–2279.

18. Djousse L, Gaziano JM. Egg consumption and risk of heart failure in the Physicians’ Health Study. Circulation 2008;117:512–516.

19. Ohlsson L, Burling H, Nilsson A. Long term effects on human plasma lipoproteins of a formulation enriched in butter milk polar lipid. Lipids Health Dis. 2009; 8:44.

20. Jain MK, Rogers J, Berg O, Gelb MH. Interfacial catalysis by phospholipase A2: activation by substrate replenishment. Biochemistry. 1991; 30:7340–7348.

21. Dietschy JM, Wilson JD. Regulation of cholesterol metabolism. 3. The N Engl J Med. 1970; 282:1241–1249.

22. Nissen SE, Tuzcu EM, Schoenhagen P, Crowe T, Sasiela WJ, Tsai J, et al. Reversal of Atherosclerosis with Aggressive Lipid Lowering I: Statin therapy, LDL cholesterol, C-reactive protein, and coronary artery disease. N Engl J Med. 2005; 352:29–38.

23. Bailey CJ, Day C. Traditional plant medicines as treatments for diabetes. Diabetes care 1989; 12:553–564.

24. Gushiken LF, Beserra FP, Rozza AL, Bérgamo PL, Bérgamo DA, Pellizzon CH. Chemical and biological aspects of extracts from medicinal plants with antidiabetic effects. Rev Diabet Stud. 2016; 13:96–112.

25. Tag H, Kalita P, Dwivedi P, Das AK, Namsa ND. Herbal medicines used in the treatment of diabetes mellitus in Arunachal Himalaya, northeast, India. J Ethnopharmacol. 2012; 141:786–795.

26. Bhamra SK, Slater A, Howard C, Johnson M, Heinrich M. The use of traditional herbal medicines amongst South Asian diasporic communities in the UK. Phytother Res. 2017; 31:1786–1794.

27. Qian F, Korat AA, Malik V, Hu FB. Metabolic effects of monounsaturated fatty acid-enriched diets compared with carbohydrate or polyunsaturated fatty acid-enriched diets in patients with type 2 diabetes: a systematic review and meta-analysis of randomized controlled trials. Diabetes care 2016; 39:1448–1457.

28. Biswas R, Chanda J, Kar A, Mukherjee PK. Tyrosinase inhibitory mechanism of betulinic acid from Dillenia indica. Food Chem. 2017; 232:689–696.

29. Biswas R, Chanda J, Kar A, Mukherjee PK. Tyrosinase inhibitory mechanism of betulinic acid from Dillenia indica. Food Chem. 2017; 232:689–696.

30. Kumar S, Kumar V, Prakash O. Enzymes inhibition and antidiabetic effect of isolated constituents from Dillenia indica. Biomed Res Int. 2013; 2013:382063.

31. Saiful YL, Armania N. Dillenia species: A review of the traditional uses, active constituents and pharmacological properties from pre-clinical studies. Pharm Biol. 2014; 52:890–897.

32. Bolzan AD, Bianchi MS. Genotoxicity of streptozotocin. Mutat Res. 2002; 512:121–134.

33. Bonner-Weir S, Trent DF, Honey RN, Weir GC. Responses of neonatal rat islets to streptozotocin: limited B-cell regeneration and hyperglycemia. Diabetes 1981; 30:64–69.

34. Ariza L, Zaguirre M, García M, Blasco E, Rabanal RM, Bosch A, Otaegui PJ. Hyperglycemia and hepatic tumors in ICR mice neonatally injected with streptozotocin. Lab Anim 2014; 43:242–249.

35. Basit A, Hydrie MZ, Ahmed K, Hakeem R. Prevalence of diabetes, impaired fasting glucose and associated risk factors in a rural area of Baluchistan province according to new ADA criteria. J Pak Med Assoc. 2002; 52(8): 357–60.

36. Kratzsch J, Ackermann W, Keilacker H, Besch W, Keller E. A sensitive sandwich enzyme immunoassay for measurement of insulin on microtiter plates. Exp Clin Endocrinol. 1990; 95(2):229–36.

37. Van DVJ. Two methods for the determination of glycogen in liver. Biochem J. 1954; 57(3):410–416.

38. Feldman AT, Wolfe D. Tissue processing and hematoxylin and eosin staining. Methods Mol Biol. 2014; 1180:31–43.

39. Salmabadi Z, Mohseni KH, Parivar K, Karimzadeh L. Effect of Grape Seed Extract on Lipid Profile and Expression of Interleukin-6 in Polycystic Ovarian Syndrome Wistar Rat Model. Int J Fertil Steril. 2017; 11:176–183.

40. Moniruzzaman M, Rokeya B, Ahmed S, Bhowmik A, Khalil MI, Gan SH. In vitro antioxidant effects of Aloe barbadensis Miller extracts and the potential role of these extracts as antidiabetic and antilipidemic agents on streptozotocin-induced type 2 diabetic model rats. Molecules. 2012; 17:12851–12867.

41. Tadin SM, Peterson LB, Cumiskey AM, Rosa RL, Mendoza VH, Castro PJ, et al. siRNA-induced liver ApoB knockdown lowers serum LDL-cholesterol in a mouse model with human-like serum lipids. J Lipid Res. 2011; 52:1084–1097.

42. Rorsman P, Ashcroft FM. Pancreatic beta-cell electrical activity and insulin secretion: of mice and men. Physiol Rev. 2018, 98:117–214.

43. Yang Q, Vijayakumar A, Kahn BB. Metabolites as regulators of insulin sensitivity and metabolism. Nat Rev Mol Cell Biol. 2018, 19:654–672.

44. Zhang X, Jin Y, Wu Y, Zhang C, Jin D, Zheng Q, Li Y. Anti-hyperglycemic and anti-hyperlipidemia effects of the alkaloid-rich extract from barks of Litsea glutinosa in ob/ob mice. Sci Rep. 2018; 8(1):12646.

45. Thorens B. GLUT2, glucose sensing and glucose homeostasis. Diabetologia 2014; 58:221–232.

46. Thorens B, Weir GC, Leahy JL, Lodish HF, Bonner WS. Reduced expression of the liver/beta-cell glucose transporter isoform in glucose-insensitive pancreatic beta cells of diabetic rats. Proc Natl Acad Sci USA 1990; 87: 6492–6496.

47. Gonzalez JA, Jensen LT, Fugger L, Burdakov D. Metabolism-Independent Sugar Sensing in Central Orexin Neurons. Diabetes 2008; 57:2569–2576.

48. Lamy CM, Sanno H, Labouèbe G, Picard A, Magnan C, Chatton JY, Thorens B. Hypoglycemia-Activated GLUT2 Neurons of the Nucleus Tractus Solitarius Stimulate Vagal Activity and Glucagon Secretion. Cell Metab 2014; 19:527–538.

49. Fu C, Yang D, Peh WY, Lai S, Feng X, Yang H. Structure and antioxidant activities of proanthocyanidins from elephant apple (Dillenia indica Linn.). J Food Sci. 2015; 80:2191–2199.

50. Olennikov DN, Chirikova NK, Kashchenko NI, Nikolaev VM, Kim SW, Vennos C. Bioactive phenolics of the genus Artemisia (Asteraceae): HPLC-DAD-ESI-TQ-MS/MS Profile of the Siberian species and their inhibitory potential against alpha-amylase and alpha-glucosidase. Front Pharmacol. 2018; 9:756.

51. Khan MF, Rawat AK, Khatoon S, Hussain MK, Mishra A, Negi DS. In vitro and in vivo antidiabetic effect of extracts of Melia azedarach, Zanthoxylum alatum, and Tanacetum nubigenum. Integr Med Res. 2018; 7(2):176–183.

52. Ota A, Ulrih NP. An overview of herbal products and secondary metabolites used for management of type two diabetes. Front Pharmacol. 2017; 84:36.

53. Chang SK, Kohlgruber AC, Mizoguchi F, Michelet X, Wolf BJ, Wei K, Lee PY, Lynch L, Duquette D, Ceperuelo MV, et al. Stromal cell cadherin-11 regulates adipose tissue inflammation and diabetes. J Clin Invest 2017; 127(9): 3300–3312.

54. Jones BJ, Tan T, Bloom SR. Minireview: Glucagon in stress and energy homeostasis. Endocrinology 2012; 153(3):1049–54.

55. Chen Q, Mo R, Wu N, Zou X, Shi C, Gong J, Li J, Fang K, Wang D, Yang D, et al. Berberine ameliorates diabetes-associated cognitive decline through modulation of aberrant inflammation response and insulin signaling pathway in DM rats. Front Pharmacol. 2017; 8:334.

56. Gadi R, Samaha FF. Dyslipidemia in type 2 diabetes mellitus. Curr Diab Rep. 2007; 7(3):228–234.

57. Xie ZQ, Liang G, Zhang L, Wang Q, Qu Y, Gao Y, Lin LB, Ye S, Zhang J, Wang H, et al. Molecular mechanisms underlying the cholesterol-lowering effect of Ginkgo biloba extract in hepatocytes: a comparative study with lovastatin. Acta Pharmacol Sin. 2009; 30:1262–1275.

58. Desmarchelier C, Ludwig T, Scheundel R, Rink N, Bader BL, Klingenspor M, et al. Diet-induced obesity in ad libitum-fed mice: food texture overrides the effect of macronutrient composition. Br J Nutr. 2013;109:1518–1527.

59. Woods SC, Seeley RJ, Rushing PA, D’Alessio D, Tso P. A controlled high-fat diet induces an obese syndrome in rats. J Nutr. 2003;133:1081–1087.

60. Nascimento AF, Sugizaki MM, Leopoldo AS, Lima LAP, Luvizotto RA, Nogueira CR, et al. A hypercaloric pellet-diet cycle induces obesity and co-morbidities in Wistar rats. Arq Bras Endocrinol Metabol. 2008; 52:968–974.

61. Matheus VA, Monteiro L, Oliveira RB, Maschio DA, Collares BCB, Butyrate reduces high-fat diet-induced metabolic alterations, hepatic steatosis and pancreatic beta cell and intestinal barrier dysfunctions in prediabetic mice. Exp Biol Med. 2017; 242:1214–1226.

62. Grundy SM. Does Dietary Cholesterol Matter? Curr Atheroscler Rep 2016; 18:68.

63. Bunzel B. Psychological problems and management of cancer patients from the viewpoint of the clinical psychologist. Wien Med Wochenschr. 1989; 139:498–500.

64. Muthuramu I, Amin R, Postnov A, Mishra M, Jacobs F, Gheysens O, et al. Coconut Oil Aggravates Pressure Overload-Induced Cardiomyopathy without Inducing Obesity, Systemic Insulin Resistance, or Cardiac Steatosis. Int J Mol Sci. 2017; 18.

65. Son J, Grant RJ, Larson LL. Effects of tallow and escape protein on lactational and reproductive performance of dairy cows. J Dairy Sci. 1996; 79:822–830.

66. Rashid KM, Ahsan H, Siddiqui S, Siddiqui WA. Tocotrienols have a nephroprotective action against lipid-induced chronic renal dysfunction in rats. Ren Fail. 2015; 37:136–143.

67. Matsuo T, Shimomura Y, Saitoh S, Tokuyama K, Takeuchi H, Suzuki M. Sympathetic activity is lower in rats fed a beef tallow diet than in rats fed a safflower oil diet. Metabolism. 1995; 44:934–939.

68. Pascual FV. Usefulness of plant sterols in the treatment of hypercholesterolemia. Nutr Hosp. 2017; 34:62–67.

69. Li Y, Qin G, Liu J, Mao L, Zhang Z, Shang J. Adipose tissue regulates hepatic cholesterol metabolism via adiponectin. Life Sci. 2014; 118(1):27–33.

70. Ros E. Intestinal absorption of triglyceride and cholesterol. Dietary and pharmacological inhibition to reduce cardiovascular risk. Atherosclerosis 2000; 151(2):357–379.

71. Ma Y, Li Y, Chiriboga DE, Olendzki BC, Hebert JR, Li W, Leung K, Hafner AR, Ockene IS. Association between carbohydrate intake and serum lipids. J Am Coll Nutr. 2006; 25(2):155–163.

72. Kumar S, Kumar V, Prakash OM. Microscopic evaluation and physiochemical analysis of Dillenia indica leaf. Asian Pac J Trop Biomed. 2011; 1(5):337–340.

73. Burg VK, Grimm HS, Rothhaar TL, Grosgen S, Hundsdorfer B, Haupenthal VJ, et al. Plant sterols the better cholesterol in Alzheimer’s disease? A mechanistical study. J Neurosci. 2013; 33:16072–16087.

74. Çimen I, Kocatürk B, Koyuncu S, Tufanlı Ö, Onat UI, Yıldırım AD, et al. Prevention of atherosclerosis by bioactive palmitoleate through suppression of organelle stress and inflammasome activation. Sci Transl Med. 2016; 8:358ra126–358ra126.

75. Somani SJ, Badgujar LB, Sutariya BK, Saraf MN. Protective effect of Dillenia indica L. on acetic acid induced colitis in mice. Indian J Exp Biol. 2014; 52:876–881.

76. Nguyen TT. The cholesterol-lowering action of plant stanol esters. J Nutr 1999; 129(12):2109–2112.

77. De SE, Mensink RP, Plat J. Effects of plant sterols and stanols on intestinal cholesterol metabolism: suggested mechanisms from past to present. Mol Nutr Food Res. 2012; 56(7):1058–1072.

78. Hunter PM, Hegele RA. Functional foods and dietary supplements for the management of dyslipidaemia. Nat Rev Endocrinol. 2017; 13(5):278–288.

79. Trautwein EA, Koppenol WP, de JA, Hiemstra H, Vermeer MA, Noakes M, Luscombe MND. Plant sterols lower LDL-cholesterol and triglycerides in dyslipidemic individuals with or at risk of developing type 2 diabetes; a randomized, double-blind, placebo-controlled study. Nutr Diabetes. 2018; 8(1):30.

80. Mineo S, Noguchi A, Nagakura Y, Kobori K, Ohta T, Sakaguchi E, et al. Boysenberry Polyphenols Suppressed Elevation of Plasma Triglyceride Levels in Rats. J Nutr Sci Vitaminol. 2015; 61:306–312.

81. Nishibori N, Kishibuchi R, Morita K. Suppressive effect of Okara on intestinal lipid digestion and absorption in mice ingesting high-fat diet. Int J Food Sci Nutr 2018; 69:690–695.

82. McLaren DG, Han S, Murphy BA, Wilsie L, Stout SJ, Zhou H, et al. DGAT2 Inhibition Alters Aspects of Triglyceride Metabolism in Rodents but Not in Non-human Primates. Cell Metab. 2018; 27:1236–1248.e1236.

83. Kuivenhoven JA, Groen AK. Beyond the genetics of HDL: why is HDL cholesterol inversely related to cardiovascular disease? Handb Exp Pharmacol. 2015; 224:285–300.

84. Yvan CL, Ranalletta M, Wang N, Han S, Terasaka N, Li R, et al. Combined deficiency of ABCA1 and ABCG1 promotes foam cell accumulation and accelerates atherosclerosis in mice. J Clin Invest. 2007; 117:3900–3908.

85. Bonetti PO, Lerman LO, Lerman A. Endothelial dysfunction: a marker of atherosclerotic risk. Arterioscler Thromb Vasc Biol. 2003; 23:168–175.

86. Ahotupa M. Oxidized lipoprotein lipids and atherosclerosis. Free Radic Res. 2017; 51:439–447.

87. Greaves DR, Gordon S. The macrophage scavenger receptor at 30 years of age: current knowledge and future challenges. J Lipid Res. 2009; 50 Suppl:S282–286.

88. Libby P, Ridker PM, Hansson GK. Progress and challenges in translating the biology of atherosclerosis. Nature. 2011; 473:317–325.

89. Annema W, Von EA, Kovanen PT. HDL and atherothrombotic vascular disease. Handb Exp Pharmacol. 2015; 224:369–403.

90. Drechsler M, Megens RT, Van ZM, Weber C, Soehnlein O. Hyperlipidemia-triggered neutrophilia promotes early atherosclerosis. Circulation. 2010; 122:1837–1845.

91. Serbina NV, Pamer EG. Monocyte emigration from bone marrow during bacterial infection requires signals mediated by chemokine receptor CCR2. Nat Immunol. 2006; 7:311–317.

92. Shi C, Jia T, Mendez FS, Hohl TM, Serbina NV, Lipuma L, et al. Bone marrow mesenchymal stem and progenitor cells induce monocyte emigration in response to circulating toll-like receptor ligands. Immunity. 2011; 34:590–601.

93. Brownlee M. Biochemistry and molecular cell biology of diabetic complications. Nature 2001; 414(6865):813–820.

94. Nakagawa T, Tanabe K, Croker BP, Johnson RJ, Grant MB, Kosugi T, Li Q. Endothelial dysfunction as a potential contributor in diabetic nephropathy. Nat Rev Nephrol. 2011; 7(1):36–44.

95. Fraser A, Harris R, Sattar N, Ebrahim S, Davey SG, Lawlor DA. Alanine aminotransferase, -glutamyltransferase, and incident diabetes: the british women’s heart and health study and meta-analysis. Diabetes care 2009; 32(4):741–750.

96. Chen SC, Tsai SP, Jhao JY, Jiang WK, Tsao CK, Chang LY. Liver fat, hepatic enzymes, alkaline phosphatase and the risk of incident type 2 diabetes: a prospective study of 132,377 adults. Sci Rep. 2017; 7(1):4649.

97. Kewcharoenwong C, Rinchai D, Nithichanon A, Bancroft GJ, Ato M, Lertmemongkolchai G. Glibenclamide impairs responses of neutrophils against Burkholderia pseudomallei by reduction of intracellular glutathione. Sci Rep. 2016; 6: 34794.

